# Predicting barrier architecture and the genomic landscape of differentiation under polygenic divergent selection

**DOI:** 10.64898/2026.09.03.749118

**Authors:** Arthur Zwaenepoel

## Abstract

We consider polygenic divergent selection in a mainland-island model, where our aim is to understand how patterns of genetic variation along the genome reflect the genetic architecture of postzygotic reproductive isolation. We derive a new expression for the effective migration rate (*m*_*e*_) at both neutral and divergently selected loci (i.e. barrier loci), and develop a numerical approach to co-predict *m*_*e*_ and allele frequencies at barrier loci. Using this *m*_*e*_, we predict neutral coalescence times along the genome, and show how our results for the mainland-island model can be used to obtain coalescence time predictions for nonequi-librium demographic models. We validate our approach extensively using forward-in-time individual-based simulations and show that our *m*_*e*_-based approximation remains accurate across a broad parameter range. We study both the case of a single barrier locus and a pair of barrier loci in detail to identify the conditions under which an *m*_*e*_-based approach performs well. We use our new predictions to examine how the genetic architecture of divergent selection shapes barriers to gene flow, quantifying hitchhiking effects along the genome and evaluating the effects of polygenicity on reproductive isolation. We discuss the implications for mapping barriers to gene flow using population genomic data, both for model-based inference and genome scan approaches based on summary statistics.

## Introduction

Uncovering the genetic architecture of reproductive isolation (RI) between diverging populations is a central goal of speciation genomics (Seehausen et al., 2014; Stankowski et al., 2023). Usually, population genomic studies try to achieve this by analyzing the genomic landscape of differentiation (as for instance quantified by *F*_ST_), where it is assumed that regions of elevated differentiation could signal the presence of barriers to gene flow (Wolf and Ellegren, 2017; Ravinet et al., 2017). However, the interpretation of such patterns of genomic differentiation has been troubled by the lack of good *a priori* theoretical predictions (Cruickshank and Hahn, 2014; Wolf and Ellegren, 2017; Lohse, 2017). Indeed, predicting observable genetic variation within and between populations when selection acts on many loci remains a core challenge of (theoretical) population genetics. Solid theoretical results are largely restricted to single-locus and two-locus models, while detailed multilocus predictions remain, in general, analytically intractable even in the large population limit where populations evolve deterministically, let alone for small populations where genetic drift cannot be ignored.

That said, a number of fruitful approaches have emerged that make use of suitably defined *effective* parameters. The latter enable the approximation of a complicated multilocus equilibrium by means of single-locus population genetic theory. For instance, the effects on neutral variation of purifying selection at numerous targets in the genome can often be modeled by means of an *effective population size* (*N*_*e*_) that varies along the genome (Hudson and Kaplan, 1995; Nordborg et al., 1996; Cvijović et al., 2018; Santiago and Caballero, 2016; Buffalo and Kern, 2024). Similarly, the effects of polygenic divergent selection between populations have been succesfully modeled by means of an *effective migration rate* that varies along the genome (Barton and Bengtsson, 1986; Juric et al., 2016; Aeschbacher and Bürger, 2014; Aeschbacher et al., 2017; Sachdeva, 2022; Zwaenepoel et al., 2024). While powerful, these approaches depend on a number of assumptions and usually break down outside of certain parameter regimes. Furthermore, it is not always clear whether in those regimes an effective parameter actually ‘exists’, i.e. whether single-locus theory can adequately capture what happens at any individual locus in a multilocus system. This is not merely a theoretical question, but bears on the interpretation of model-based inference methods for uncovering barriers to gene flow (e.g. Laetsch et al. (2022), Fraïsse et al. (2021), Burban et al. (2024)). Such methods infer effective parameters that vary along the genome, but what exactly this is capturing and how it relates to the genetic architecture of RI remains rather unclear.

Here we consider multilocus divergent selection in a haploid mainland-island model at equilibrium. In the context of this model, our goals are twofold: (1) predict the genomic landscape of *neutral* genetic variation within and between populations and (2) predict allele frequency divergence between mainland and island at *selected* loci. Our approach is to build an approximation for multilocus migration-selection-drift equilibrium using single-locus theory, together with a suitably defined effective migration rate *m*_*e*_ = *m × g*, where *g*, the local reduction in gene flow, is called the *gene flow factor* (gff) (Bengtsson, 1985) and is assumed to vary along the genome. Below, we present a new *m*_*e*_ approximation which improves significantly on previous approaches. Specifically, we obtain accurate predictions of both neutral differentiation and adaptive divergence even when selection is weak relative to migration and/or genetic drift (unlike Aeschbacher and Bürger (2014)), and when linkage among barrier loci is tight (unlike Sachdeva (2022); Zwaenepoel et al. (2024)).

We distinguish two kinds of *m*_*e*_: (1) neutral *m*_*e*_ and (2) selective *m*_*e*_ (see also Kobayashi et al. (2008)). Neutral *m*_*e*_ quantifies the rate at which lineages at a neutral locus move between populations at equilibrium. This can be considered both forward and backward in time. Backward in time, the neutral *m*_*e*_ in the mainland-island model corresponds to the rate at which a neutral lineage sampled from the island population is absorbed into the mainland. Denoting the waiting time until absorption (at equilibrium) by *T* , we will have *m*_*e*_ = 1*/*E[*T* ]. Considered forward in time, *m*_*e*_ is the rate at which a migrant allele at a neutral locus is transferred onto the resident genetic background. When migration is not too strong, this rate is determined by the *reproductive value* (RV) of a migrant individual, i.e. the expected long term contribution of a migrant to the local gene pool (Kobayashi et al., 2008; Rousset, 2004). It is currently unclear under what conditions a neutral *m*_*e*_ suffices to statistically predict genealogies at neutral loci linked to a set of barrier loci. This question is becoming increasingly relevant with the recent growth of ancestral recombination graph (ARG) inference approaches and methods that seek to learn about gene flow and natural selection from inferred genealogies along the genome (Nielsen et al., 2025).

Neither the forward nor the backward time formulation works for gene flow at loci that are themselves subject to divergent selection and are polymorphic on the island at migration-selection-drift equilibrium (see below). Nevertheless, we would still like to use an effective migration rate to account for the effects of multilocus selection on equlibrium allele frequencies. The selective *m*_*e*_, then, is defined as the migration rate which would yield the observed equilibrium allele frequency divergence if there were no other selected loci. Phrased differently, it is the migration rate that, when “plugged into” single-locus theory, yields the equilibrium allele frequency divergence at the selected locus when in a multilocus system. As with neutral *m*_*e*_, it is currently unclear to what extent a selective *m*_*e*_ really exists: can the marginal allele frequency distribution at any selected locus at equilibrium be described by the single locus equilibrium distribution with a rescaled *m*? For weak linkage, Sachdeva (2022) and Zwaenepoel et al. (2024) derived *m*_*e*_ approximations that do enable accurate prediction of equilibrium allele frequencies in a mainland-island and infinite-island model, and they showed that, in this regime, the selective *m*_*e*_ is essentially the same as the neutral *m*_*e*_. With more tight linkage (*r < s*), the approximations of the latter authors break down, but this does not mean that there is no *m*_*e*_ that would do the job.

Using our novel *m*_*e*_ approximation together with phase-type theory (Hobolth et al., 2024), we are able to predict the genomic landscape of neutral differentiation for a rather general class of demographic models, and using forward simulations we show that these predictions are remarkably accurate. To our knowledge, such accurate and detailed predictions of the effects of multilocus divergent selection on neutral variation along the genome are unprecedented, paving the way for improved model-based genomic inference of barriers to gene flow. We use the *m*_*e*_ approximation to gain more insights in how the genetic architecture of divergent selection (i.e. the number of selected loci, their genomic distribution and effect sizes) shapes the architecture of RI (the barrier strength along the genome). Specifically, we show how the selective *m*_*e*_ can be used to quantify the extent of divergence hitchhiking, where divergently selected allels are protected from swamping due to closely linked barrier loci, relative to genome hitchhiking, where alleles are protected from swamping due to the genome-wide barrier effect (Nosil, 2012). We further use our new *m*_*e*_ predictions to investigate the conditions under which polygenicity and the spatial distribution of selected loci weaken or strengthen the barrier to gene flow. In the discussion section we consider the implications for empirical speciation genomics, focusing in particular on the mapping of barrier loci from population genomic data.

## Model and Methods

### Multilocus mainland-island model

We consider a pair of haploid Wright-Fisher populations, population *A* (the mainland) of size *N*_*A*_ and population *B* (the island) of size *N*_*B*_ (see table 1 for a glossary of the relevant notation). In each generation a random number *M* of individuals on the island are replaced by migrants from the mainland population, where *M* ∼ Poisson(*mN*_*B*_). Fitness on the island is determined by *L* biallelic loci with selection coefficients *s*_1_, *s*_2_, … , *s*_*L*_. The *L* loci are located at map positions *x*_1_, *x*_2_, … , *x*_*L*_, spread along a single chromosome of map length *C* Morgan. We write *r*_*ij*_ for the recombination rate between selected locus *i* and *j*, and *r*_*i*_ (or more explicitly *r*_*i*_(*x*)) for the recombination rate between a generic neutral locus at position *x* and locus *i*. We assume fitness effects combine multiplicatively, i.e. let *X*_*ij*_ ∈ {0, 1} be the allelic state at locus *j* in individual *i*, then individual *i* has fitness 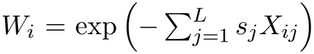 on the island. We write *q*_*i*_ = 1 *− p*_*i*_ for the frequency of the locally deleterious allele (`1`) on the island at locus *i*. Throughout, we assume the mainland is fixed for the `1` allele at each selected locus, but this assumption is not essential. We assume mutations occur at rate *u* ≪ *s*_*i*_ on the island, irrespective of allelic state.

**Table 1.** glossary for the notation used in the main text.

| **notation** | **description** |
|---|---|
| *N*<sub>*A*</sub> | mainland population size |
| *N*<sub>*B*</sub> | island population size |
| *m* | migration rate |
| *L* | number of selected loci |
| *s*<sub>*i*</sub> | selection coefficient at locus *i* |
| *x*<sub>*i*</sub> | map position of selected locus *i* |
| *r*<sub>*i*</sub> | recombination rate between a focal neutral locus and barrier locus *i* |
| *r*<sub>*ij*</sub> | recombination rate between barrier loci *i* and *j* |
| *m*<sub>*e*</sub>/*m*<sub>*e*</sub>(*x*) | neutral effective migration rate at a focal neutral locus/at genomic position *x* |
| *g*/*g*(*x*) | *ibid.* gene flow factor (gff) |
| *m*<sub>*e*,*i*</sub>/*g*<sub>*i*</sub> | effective migration rate/gff at barrier locus *i* |
| *b* | barrier strength (*b* = *g*<sup>−1</sup>) |
| *t*<sub>*A*</sub> | coalescence time of a pair of lineages in population *A* |
| *t*<sub>*AB*</sub> | coalescence time for a pair of lineages from populations *A* and *B* |
| *t*<sup>\*</sup><sub>.</sub> | neutral prediction for 𝔼[*t*<sub>.</sub>] with *m*<sub>*e*</sub> substituted for *m* |
| *t*<sub>*d*</sub> | divergence (split) time of populations *A* and *B* |
| *h* | proportion of recent migrant ancestry (used in mixture approximation) |

### Neutral effective migration rate

We first develop an approximation for the effective migration rate at neutral loci, assuming allele frequencies at selected loci (*p*_*i*_) are known. Later we will consider the *m*_*e*_ at *selected* loci and outline a numerical approach for predicting the associated allele frequencies. Our derivation is based on the correspondence between reproductive value (RV) and *m*_*e*_ (Kobayashi et al., 2008; *Sachdeva, 2022; Westram et al*., *2022), but where we consider the RV from the “gene’s eye view”. Specifically, we use a multi-type branching process (BP) to approximate the RV of a migrant gene* entering the resident island population at equilibrium. Crucially, our approach assumes that *m* ≪ 1, so that the gff is well approximated by the RV, but *not* that *m* ≪ *s* (as e.g. the approximation by Aeschbacher and Bürger (2014) does).

#### A single barrier locus

To illustrate our approach, we first analyze the case of a neutral locus associated with a single barrier locus. The BP has two types, corresponding to distinct descendants of a single migrant individual. Type 1 individuals correspond to descendants which have inherited both the neutral and selected allele from the initial migrant, whereas type 2 individuals have inherited only the neutral allele. It should be stressed that the types do *not* correspond to particular genotypes, but to different kinds of descendants from an initial migrant individual, whose genotypes will depend on the composition of both the migrant and resident gene pool.

The resident population is assumed to be at migration-selection equilibrium, with deleterious allele frequency *q*. The fitness of a type 1 individual, relative to the mean fitness of the resident population, is

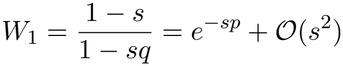

When migration is sufficiently weak, we can ignore the possibility that migrants and their descendants mate with each other. A type 1 individual will then only mate with resident individuals. When a recombination event occurs between the two loci, the offspring will be of type 2, in which case the allele at the barrier locus will be a random draw from the resident gene pool. The *expected* relative fitness of a type 2 individual is hence 1.

Let *Z*_*t*,1_ and *Z*_*t*,2_ be the number of type 1 and type 2 descendants of a migrant after *t* generations of random mating on the island. The BP describing the evolution of *Z*_*t*_ = (*Z*_*t*,1_, *Z*_*t*,2_) has mean matrix

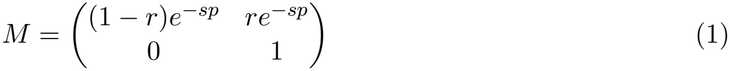

The RV of the neutral gene copy introduced by the migrant individual, which is equal to the gff *at the neutral locus* in the limit of weak migration, can be found as the expected number of times a neutral allele inherited from the migrant ‘escapes’ into the resident background, i.e.

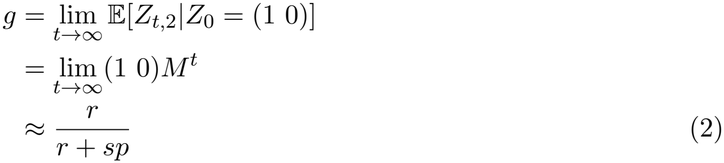

The same result is obtained by a backward-time analysis of the time to absorption into the mainland (section S1.1). Equation (2) is equivalent to equation B8 of Zwaenepoel et al. (2024) if *p* = 1 *− m/s* is assumed (i.e. deterministic migration-selection balance). When *m* ≪ *s* so that *p →* 1 (i.e. there is complete divergence between mainland and island at the selected locus), eq. (2) corresponds to the approximation of Petry (1983) (see also Bengtsson (1985); Aeschbacher and Bürger (2014)).

#### Two barrier loci

We now extend this approach to derive the gff at a neutral locus associated with *two* barrier loci, before generalizing to the *L*-locus setting. We start by considering the case where the neutral locus is on either side of the two barrier loci, and then analyze the case where the neutral locus is located in between the two barriers.

Let the neutral locus be on the left of the two selected loci (the other case follows by symmetry). The recombination rates between the neutral locus and the first and the second selected locus are *r*_1_ and *r*_2_ respectively. The recombination rate between the two selected loci is *r*. We assume *r ≈ r*_2_ *−r*_1_, i.e. recombination rates are small enough that we can ignore multiple crossovers. The BP now has three types, corresponding to distinct kinds of descendants of a migrant individual that have inherited the focal neutral allele from their migrant ancestor. Specifically, we track offspring where the introgressing neutral allele is associated with (1) migrant alleles at both selected loci, (2) the migrant allele at the leftmost selected locus and (3) neither of the migrant alleles. We can again write down a mean matrix for the BP:

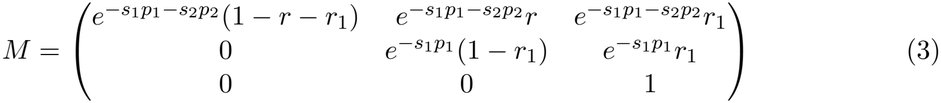

A similar analysis as the one leading to eq. (2) yields

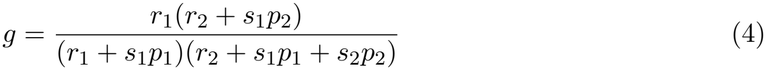

to leading order in *r* and *s* parameters. For the case where the neutral locus is in between the two selected loci, we can again write down the mean matrix of the associated BP (which now has four types), and determine the gff accordingly, yielding

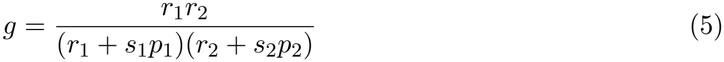

Again, in the limit of complete divergence (when *m/s*_*i*_ *→* 0, so that *p*_*i*_ *→* 1), these expressions are identical to those in Aeschbacher and Bürger (2014).

#### Multilocus generalization

These results can be generalized to a chromosome with *L* selected loci. Consider a focal neutral locus at map position *x*, and let *L*_*l*_ (*L*_*r*_) be the set of selected loci on the left (right) of the focal locus. We fix an ordering of the loci in these sets from closer to further removed from the focal locus, i.e. for *i, j* ∈ *L*_*l*_, *i < j* if |*x*_*i*_ *− x*| *<* |*x*_*j*_ *− x*|. The gff at the neutral locus can be expressed as

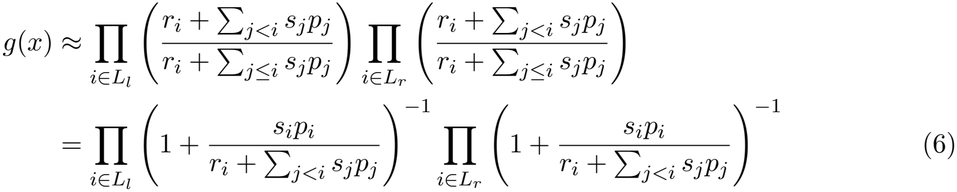

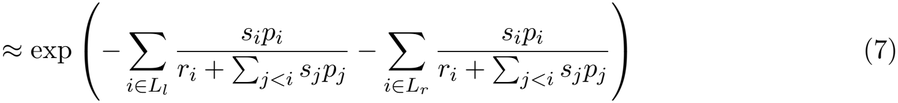

Equation (6) is written in the same form as Aeschbacher and Bürger (2014) (their equation 24), highlighting again that the the effective migration rate predicted by this approach is equivalent to the latter but with *s*_*i*_*p*_*i*_ substituted for *s*_*i*_ (see also Westram et al. (2022), SI2 eq(5)). Further-more, for loose linkage (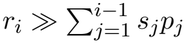 for all *i*), eq. (7) is equivalent to the gff of Zwaenepoel et al. (2024), restricted to the haploid setting. Hence, for *unlinked* loci, the theory of the latter authors applies, so that when there are multiple chromosomes, each unlinked locus *j* further contributes a factor 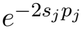 to the gff.

### Predicting allele frequencies at selected loci

#### Selective *m*_*e*_

The effective migration rate for a selected locus can not be defined in the same way as for a neutral locus: deleterious alleles introduced by migration are purged eventually, and the long-term contribution to the island gene pool vanishes. Indeed, if we consider the neutral *m*_*e*_ derived above (eq. (7)) and take the limit as the recombination rate to the closest selected locus becomes zero, we find *g* = 0 and hence *m*_*e*_ = 0. Using this *m*_*e*_ to predict equilibrium allele frequency divergence would hence imply complete divergence between mainland and island. When linkage is sufficiently weak (*r/s >* 2, roughly), one can predict allele frequencies at selected loci reasonably well by substituting the neutral *m*_*e*_ (i.e. *m*_*e*_ for a hypothetical neutral locus at the same genomic location) for *m* in the diffusion approximation (Zwaenepoel et al., 2024). However, this approach breaks down for tight linkage.

We can obtain some insight from the case with two selected loci. If one takes the deterministic equilibrium predictions of Bürger and Akerman (2011) to find the migration rate *m*_*e*,1_ that would yield the equilibrium allele frequency at the first locus 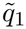 when plugged into single-locus theory (recall that 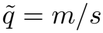 for the single-locus mainland-island model), we find

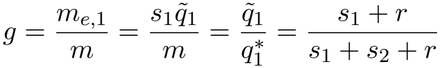

to leading order. Here 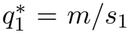 is the deterministic single-locus allele frequency prediction. Importantly, we see that the selective *m*_*e*_ is not independent of the selection coefficient *s*_1_ of the focal locus. Note that as *s*_1_ *→* 0, we obtain the neutral *m*_*e*_ for a locus linked to a single barrier locus.

We adopt a similar definition for the general case (i.e. a polygenic barrier and finite island population). At selected locus *i* ∈ [1..*L*], the gff and effective migration rate are defined as *g*_*i*_ = 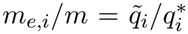 . Similar definitions have been used by other authors (Barton and Bengtsson, 1986; Kobayashi et al., 2008). Analytical predictions for the equilibrium frequency 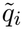 are however not available, and previous work has been restricted to the case where *m/s →* 0. We use instead our BP framework to approximate 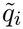 (see section S1.3). Such a BP approximation is expected to fail as *m/s* becomes large, as an introgressing deleterious allele will then no longer be sufficiently rare for the branching property to hold (i.e. their trajectories are no longer approximately independent). However, in practice we find that this approach works rather well when we use the same BP approximation for the denominator (which amounts to 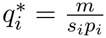).

Using this approach, and extrapolating it to the multilocus setting (see section S1.3), we find that the gff at selected locus *k* can be approximated as

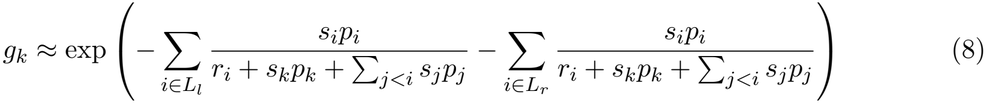

Which reduces to eq. (7) for *s*_*k*_ *→* 0. Again, when there are multiple chromosomes, every additional unlinked locus *j* contributes a factor 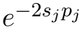 .

#### Diffusion approximation

Sachdeva (2022) showed that, given an expression for *m*_*e*_ that depends on allele frequencies, one can use an iterative approach to self-consistently predict allele frequencies at selected sites. This approach was further extended in Zwaenepoel et al. (2024) to deal with rather general genetic architectures. The idea is to predict the allele frequency at locus *j* using single-locus diffusion theory Wright (1937), substituting *m*_*e*_(*x*_*k*_) (i.e. the effective migration rate at selected locus *k*) for *m* to account for linkage disequilibrium (LD) among barrier loci. Note that *m*_*e*_(*x*_*k*_) depends on the allele frequencies at all loci, a dependence we make explicit by writing *m*_*e*_(*x*_*k*_, *p*). To predict allele frequencies at all *L* loci, we then solve the following system of *L* equations for E[*p*_*k*_], *k* ∈ [1, .., *L*]

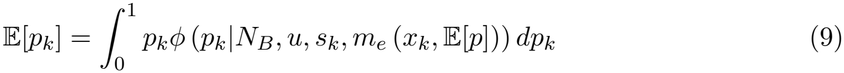

where

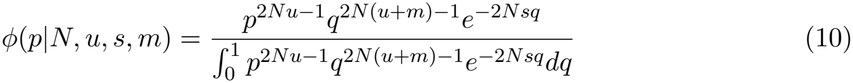

is the density of Wright’s allele frequency distribution for the haploid mainland-island model, where the mainland is fixed for the locally deleterious allele on the island.

We solve this system using a fixed point iteration. Specifically, given an initial ‘guess’ of the allele frequencies E[*p*^(0)^], we calculate *m*_*e*_ at each locus *k* as *m*_*e*_(*x*_*k*_, E[*p*^(0)^]) using eq. (8), use this *m*_*e*_ to predict E[*p*^(1)^] using eq. (9), and iterate until convergence (i.e. |E[*p*^(*k*+1)^] *−* E[*p*^(*k*)^]| *< ϵ*, for some tolerance level *ϵ*). As we have studied before (Zwaenepoel et al., 2024), this fixed point iteration may be bistable and depend on the initial state. Throughout, we assume that the initial state is complete divergence: i.e. both mainland and island are fixed for their respective locally beneficial allele frequencies. Biologically, this appears a reasonable assumption under a secondary contact scenario, when enough time has elapsed since divergence for locally beneficial alleles to reach high frequencies. We also implemented an extension of the fixed point iteration approach to deal with bidirectional migration (see section S1.4 for a detailed outline).

### Neutral coalescence times and *F*_ST_

#### Between-population coalescence time

The expected pairwise between-population coalescence time in the mainland-island model is, by definition, E[*t*_*AB*_] = 1*/m*_*e*_ + *N*_*A*_. This is just the neutral prediction, but with *m*_*e*_ substituted for *m*, which we write as 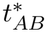 .

#### Mixture model approximation for within-population coalescence times

When sampling two lineages *within* the island, the expected coalescence time will depend on whether either of the lineages is sampled from a resident individual (i.e. an individual which has no or very distant migrant ancestry) or a recent descendant from a migrant (e.g. an F1, BC1, *etc*.) (fig. 1A). The neutral *m*_*e*_ at any genomic position quantifies the rate of gene flow from the *resident* background on the island into the mainland at that position, so, conditional on sampling two lineages that are both within a resident background, the expected coalescence time should be well predicted by 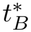 (i.e. the predicted within-island coalescence time for the neutral case, with *m*_*e*_ substituted for *m*). However, if either of the lineages is sampled within a recent descendant from a migrant, that lineage will move into the mainland on a timescale that is much shorter than 1*/m*_*e*_, and the expected coalescence time between the two island lineages will be better approximated by 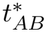, the expected between-population coalescence time for a neutral locus at the focal genomic position.

**Figure 1.**
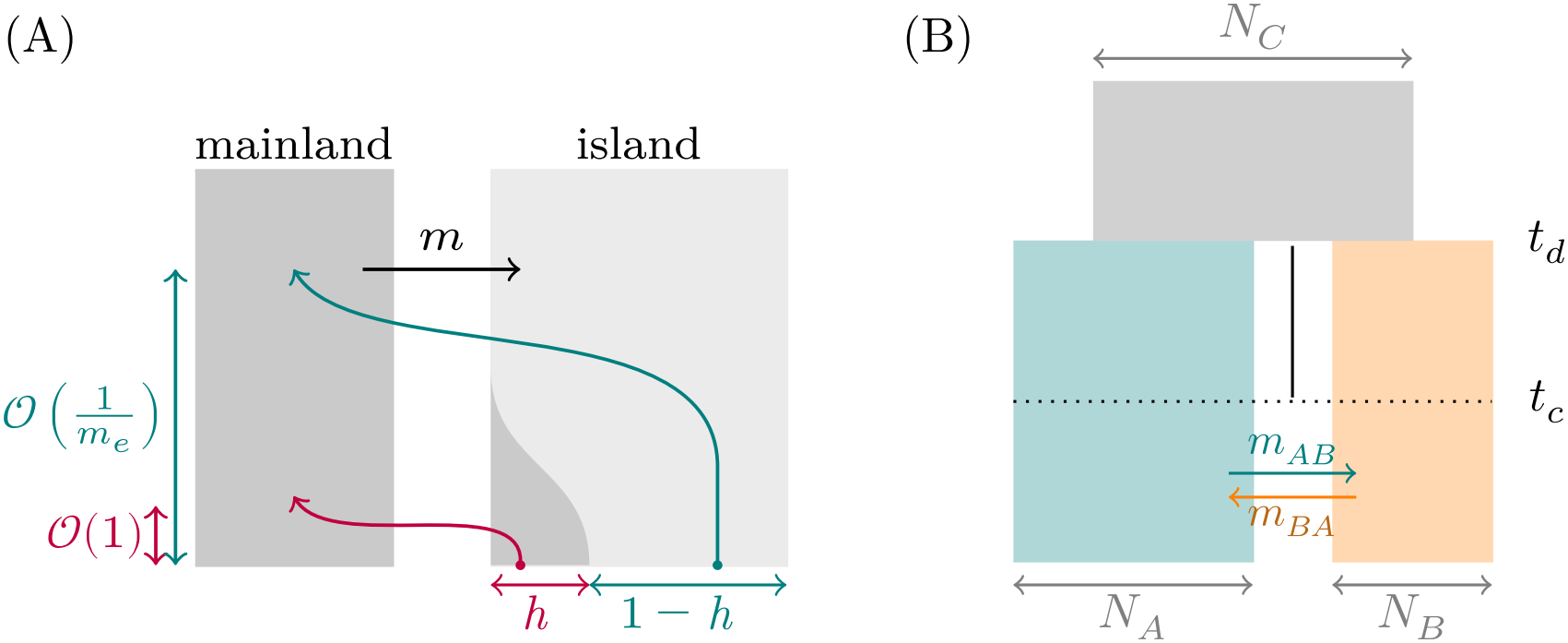
(A) Class structure within the island population results in heterogeneity of within-population coalescence times. A proportion *h* of lineages at any neutral locus traces back to the mainland on a timescale that is *O*(1) generations, while a proportion 1 *− h* finds itself on a resident background and traces back to the mainland about 1*/m*_*e*_ generations ago. With probability 2*h*(1 *− h*) *≈* 2*h*, a pair of lineages from the island coalesces on a timescale that is *O*(1*/m*_*e*_ + *N*_*A*_) and independent of the island population size. (B) General isolation-with-migration (IM) model for which we implement numerical predictions of neutral coalescence times and *F*_ST_ using phase-type theory with effective migration rates. *t*_*d*_ is the divergence time, whereas *t*_*c*_ is the time of secondary contact (during the interval [*t*_*d*_, *t*_*c*_] populations *A* and *B* are allopatric). When *t*_*d*_ = *t*_*c*_, *t*_*d*_ *→ ∞* and *m*_*BA*_ = 0, we obtain the mainland-island model that we assume in our theoretical derivations of *m*_*e*_.

This suggests that, to a first approximation, a two-component mixture model could be used to approximate the within-population coalescence time, i.e.

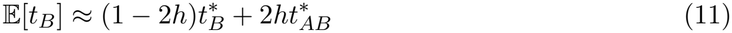

where *h* can be interpreted as the probability to sample an individual that has recent migrant ancestry (see section S1.5). We empirically tested this by fitting eq. (11) to results obtained from individual-based simulations, estimating *h* by minimizing the sum of squared deviations between predicted and observed within-population coalescence times, and we indeed found that such a two-component mixture can approximate observed within-population coalescence times very well (see section S1.5 for details).

Clearly, in theoretical predictions we cannot rely on such an *ad hoc* empirical correction, but require an *a priori* estimate for *h* given the genetic architecture of divergent selection and population genetic parameters. For the single-barrier case, the optimal mixture proportion depends on the recombination rate between the neutral locus and the barrier. When *r/s →* 0, the optimal *h* is found to be *q*, the deleterious allele frequency, supporting our heuristic interpretation where *h* is the probability of sampling an individual with recent migrant ancestry at the neutral locus. In section S1.5, we present an expression for *h* in the single-barrier model that depends on *r*, however, we find that the error introduced by using *h* = *q* at larger values of *r/s* is negligible (fig. S3). In the polygenic regime, we predict *h* using a quantitative genetics approach based one Veller et al. (2023), as outlined in detail in section S1.5. Throughout the results section we show results using the theoretically predicted *h*.

#### Isolation-with-migration demography

To obtain predictions for coalescence time distributions in more complicated demographic models, we make use of phase-type theory, employing the `PhaseGen` library (Sendrowski and Hobolth, 2026). To do so, we specify the appropriate demographic model, substituting the predicted *m*_*e*_ for *m* in the relevant epoch(s) of the demographic history. We focus in particular on variations of the isolation-with-migration model (Hey and Nielsen, 2004), depicted in fig. 1B. Throughout we assume that the onset of divergent selection coincides with the divergence time, and that by the time of secondary contact (*t*_*c*_), populations are close to fixation for their respective locally beneficial alleles (see previous section). For simplicity, we further assume for most simulations in the results section that *t*_*c*_ = *t*_*d*_ (there is no allopatric phase, and populations *A* and *B* instantaneously diverge at all divergently selected loci), that *m*_*BA*_ = 0 (unidirectional migration, so that *A* is a mainland and *B* an island population) and *N*_*C*_ = *N*_*A*_ + *N*_*B*_.

Phase-type theory allows us to numerically approximate coalescence time distributions (and hence associated moments) for arbitrary sampling configurations (Hobolth et al., 2024). Making use of the well-known correspondence between *F*_ST_ and expected coalescence times (Slatkin, 1991), we estimate *F*_ST_ between populations *A* and *B* from the pairwise coalescence times as

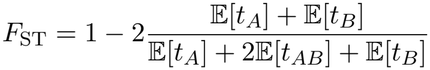

### Individual-based and coalescent simulations

We conduct forward-in-time individual-based simulations of the model specified above. We make use of tree-sequence recording (Kelleher et al., 2018) to record the ancestral recombination graph (ARG) of the population obtained at the end of the simulation. This ARG records the complete ancestry of the population along the entire genome, and allows us to query coalescence times without simulating neutral SNPs.

We make use of *recapitation* (Haller et al., 2019), simulating the ARG backward in time using `msprime` (Baumdicker et al., 2022) in those epochs of the demographic history where there is no selection. Specifically, we make use of recaptitation in two ways. Firstly, when there is unidirectional migration and the mainland has a fixed genetic composition, we use recapitation to simulate the part of the ARG that involves only mainland individuals backward-in-time. Secondly, we use recaptitation to simulate the part of the ARG in the epoch before the onset of divergent selection (which in our simulations coincides with the divergence time *t*_*d*_, but this is not necessary). It should be noted that, in the first application, recapitation implies that we can not track the full ancestry of the mainland population, as backward simulations require that the sample size *n*_*A*_ ≪ *N*_*A*_. Hence, we have to choose the number of mainland samples *n*_*A*_ that we wish to include in the simulated ARG. However, for a given forward-in-time simulation of the ‘island-part’ of the ARG, we can perform multiple replicate recapitations, and use these to estimate coalescence times.

## Results

We first investigate the efficacy of an *m*_*e*_-based approach in the context of a single barrier locus. Specifically, we consider in detail the question: when is the genealogy at a neutral locus (linked to a barrier locus) accurately approximated by a structured coalescent process with *m*_*e*_ substituted for *m*? In a similar spirit, we then look at the *selective m*_*e*_ in a two-locus system and ask: when is the equilibrium allele frequency distribution at a focal barrier locus (linked to another barrier locus) well-described by single-locus migration-selection-drift equilibrium with an effective migration rate? We then move to the polygenic regime, highlighting the power of our *m*_*e*_-based approximation for the prediction of genome-wide patterns of neutral differentiation and adaptive divergence in empirically plausible parameter regimes. We consider in some detail the effects of polygenicity and linkage on the genomic signatures left by barriers to gene flow. Finally, we briefly consider an extension of our modeling approach to accommodate bidirectional migration.

### Coalescence times near a single barrier locus

When selection is sufficiently strong relative to drift (*N*_*B*_*s >* 10, say), the expected between-population coalescence time at a neutral locus is well-predicted by 1*/m*_*e*_ + *N*_*A*_, where *m*_*e*_ is given by eq. (2) (fig. 2). When linkage is very tight (*r/s <* 0.01, say) and *m/s* appreciable, we slightly overpredict the between population coalescence time. This is because, in that case, a proportion *q* of the island population will trace back to the mainland on a much shorter timescale at the neutral locus than the predicted 1*/m*_*e*_ for resident individuals. The between-population coalescence time should then be closer to (1 *− q*)(1*/m*_*e*_) + *N*_*A*_, which indeed appears to be the case (fig. S6). Similarly, substituting *m*_*e*_ for *m* in the neutral prediction for *within*-population coalescence times yields good predictions when *r/s* is not small but breaks down as we get closer to the barrier locus (solid line in fig. 2A). In this regime, our mixture approximation appears to yield good results (dashed line in fig. 2A).

**Figure 2.**
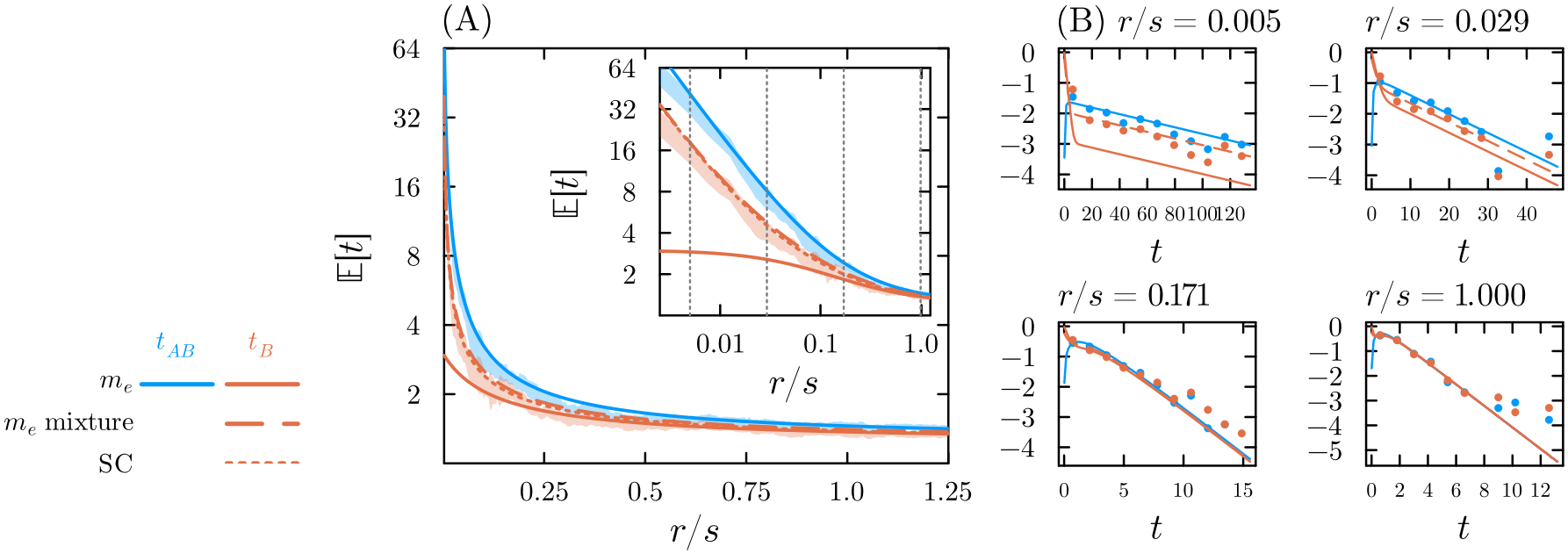
(A) Expected coalescence times (in units of *N*_*B*_ generations) at neutral loci in the presence of a single barrier to gene flow (*s* = 0.02, *m/s* = 0.2, *N*_*B*_ = 1000, *N*_*A*_ = 1000). The shaded area shows the mean *±* 2SE (standard error) for the expected between-population (*t*_*AB*_ , blue) and within-island (*t*_*B*_ , orange) coalescence time based on 100 replicate individual-based simulations. The solid and dashed lines show the neutral prediction with *m*_*e*_ substituted for *m* (eq. (2)) and the mixture approximation (eq. (11), with *h* = *q* = *m/s*) for within-population coalescence times respectively. The dotted lines are predictions from the structured coalescent (SC) model (eqs. (S8) and (S9)). Note that the structured coalescent and the *m*_*e*_ prediction are identical for *t*_*AB*_ , and that the mixture model only applies to *t*_*B*_ . (B) Distribution of pairwise coalescence times for four different map positions. The *y*-axis shows log probability density. Histograms (dots) are based on individual-based simulations, the solid lines show the phase-type distributions with *m*_*e*_ substituted for *m*, the dashed line shows the mixture approximation for the distribution of *t*_*B*_ . See also fig. S7.

Decreasing *m/s*, either by increasing the intensity of selection or decreasing the rate of migration, increases allele frequency divergence (*q*) at the barrier locus between mainland and island, increasing the time to coalescence between populations (fig. S7). The expected coalescence time within populations is also increased, but to a much lesser extent. As a result, decreasing *m/s* leads to an increase in measures of differentiation at linked neutral loci (such as *F*_ST_).

Not only does an *m*_*e*_-based approximation allow for accurate prediction of *expected values* of pairwise coalescence times, we also appear to predict the entire *distribution* of coalescence times quite well (fig. 2B). Importantly, this implies that *m*_*e*_, *N*_*A*_ and *N*_*B*_ suffice to predict *site*-based statistics (such as the within- or between-population nucleotide diversity), suggesting in turn that site-based measures *per se* do not provide much information about the genetic architecture of divergent selection beyond *m*_*e*_.

The results in fig. 2 show equilibrium features, assuming the island has been separated from the mainland and subject to migration-selection balance for an indefinite time. In the more empirically relevant scenario where the island split from the mainland *t*_*d*_ generations ago (the IM model), the signature left by a barrier locus at linked neutral sites depends strongly on *t*_*d*_ (fig. S8) (here we assume *t*_*d*_ coincides with the onset of divergent selection and gene flow, and migration-selection balance is attained instantaneously). Importantly, we find that expected coalescence times and *F*_ST_ at neutral loci in the vicinity of the barrier locus can still be accurately predicted using *m*_*e*_ (fig. S8). This implies that, while our *m*_*e*_ approximation depends crucially on the assumption of migration-selection-drift equilibrium *at selected loci*, we do not require equilibrium demography to succesfully use *m*_*e*_ to predict site-based measures at linked *neutral* loci.

### Two barrier loci

When there is a single barrier locus, the (selective) *m*_*e*_ *at* the barrier locus is, by definition, just *m* (see Methods). However, when we move to multilocus barriers, selection at any one locus affects equilibrium allele frequencies at all other loci, so that *m*_*e,i*_ *< m*. In previous work, we obtained very good predictions for equilibrium allele frequencies when barriers are polygenic and weakly linked by using a mean-field approach with a locus-specific *m*_*e*_ and the classical single-locus diffusion approximation (Zwaenepoel et al., 2024). Using a similar approach, but with the gff approximation developed in the present paper (eq. (8)), we predict allele frequencies remarkably well already in the case with only two barrier loci, even when linkage between the barrier loci becomes tight (*r/s <* 1, where *r* is the recombination rate between barrier loci, fig. 3A)^1^.

**Figure 3.**
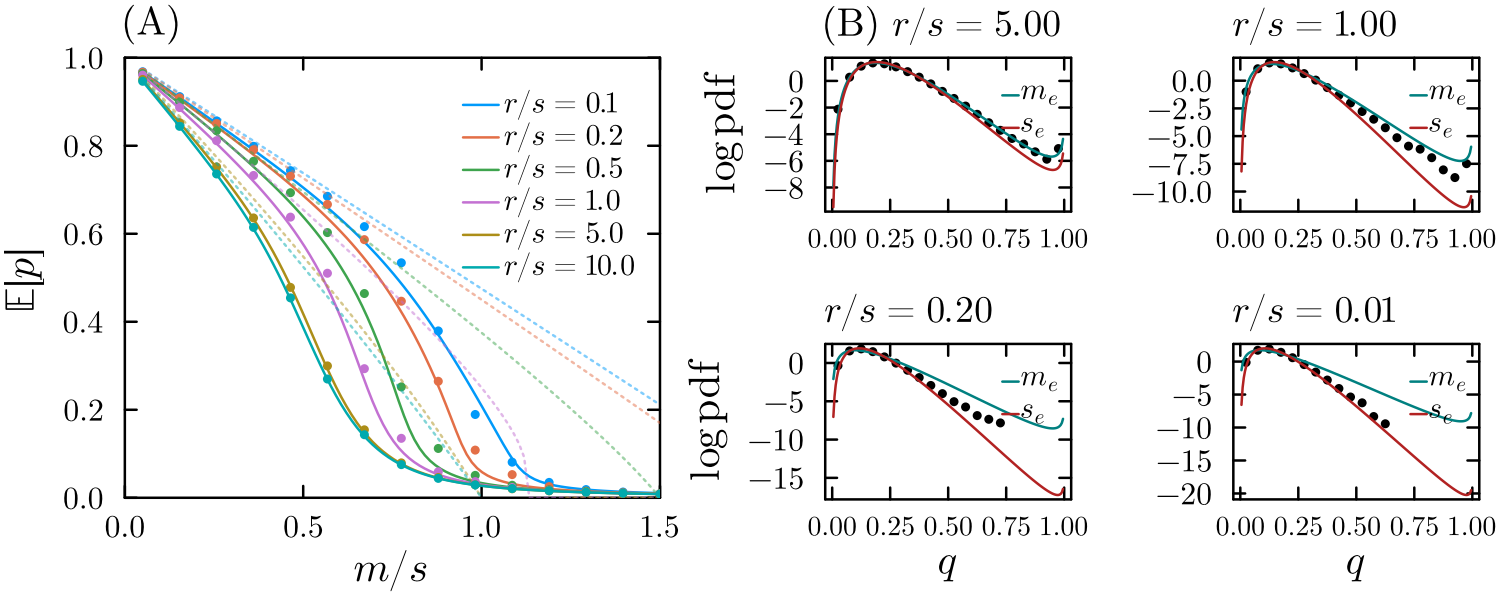
Selected allele frequencies for a two-locus barrier model. Recombination between the barrier loci occurs at rate *r*, and both loci have selection coefficient *s*. (A) Expected frequency of the locally beneficial allele on the island at the barrier loci for increasing rates of migration *m* and different recombination rates *r* between the barrier loci. The lines are theoretical predictions based on eq. (8) with the fixed point iteration, while the dots are results from individual-based simulations. The dotted lines show deterministic predictions (Bürger and Akerman, 2011). (B) Marginal equilibrium allele frequency distribution at the barrier loci for four different recombination rates, for *m/s* = 0.25. The dots show results from individual-based simulation (based on 50.000 samples). The teal line shows the prediction based on Wright’s distribution with *m*_*e*_ = *mg* substituted for *m*, while the red line shows the prediction based on Wright’s distribution with *s*_*e*_ = *s/g* substituted for *s* (where *g* is calculated according to eq. (8)). We assume *N*_*B*_*s* = 10 for both loci and *N*_*B*_ = 1000 in all simulations.

When linkage between the barrier loci is weak (*r/s >* 1), the *m*_*e*_ approximation does not only predict expected allele frequencies, but also accurately predicts the allele frequency distribution at the barrier loci (fig. 3B). However, when linkage becomes tight, the two loci start to behave essentially like a single selected locus with an increased selection coefficient (Barton, 1983). As a result, the allele frequency distribution begins to deviate from the *m*_*e*_-based prediction, and an approach based on an effective *selection coefficient* provides a better description of migration-selection equilibrium at the barrier loci (fig. 3B, red line).

For the case with two barrier loci, a backward-time (structured coalescent) analysis based on tracking the movement of a neutral lineage between different genetic backgrounds (Nordborg, 1997) remains feasible and is expected to provide an accurate description as long as *N*_*B*_*s* is sufficiently large (see section S1.2). Hence, we can compare our predictions based on eq. (7) both against individual-based simulations and numerical backward-time predictions. In fig. 4, we focus on the case where migration is strong relative to selection (here *m/s* = 0.5), which is the more challenging regime for theoretical predictions. Overall, our predictions for *between*-population coalescence times using the *m*_*e*_ given by eq. (7) appear virtually indistinguishable from those obtained using the backward-time analysis, and both predict the expected coalescence time as estimated from simulations very accurately (fig. 4). Note that for the structured coalescent predictions, we assume allele frequencies at the selected loci and LD between them as given (estimated from simulations), whereas in the *m*_*e*_-based predictions the allele frequencies are co-predicted. In contrast with the *m*_*e*_ from Aeschbacher and Bürger (2014), the approximation developed here can deal adequately with the case where there is strong migration (*m* ∼ *s*) and hence only partial divergence between mainland and island (see also fig. S9).

**Figure 4.**
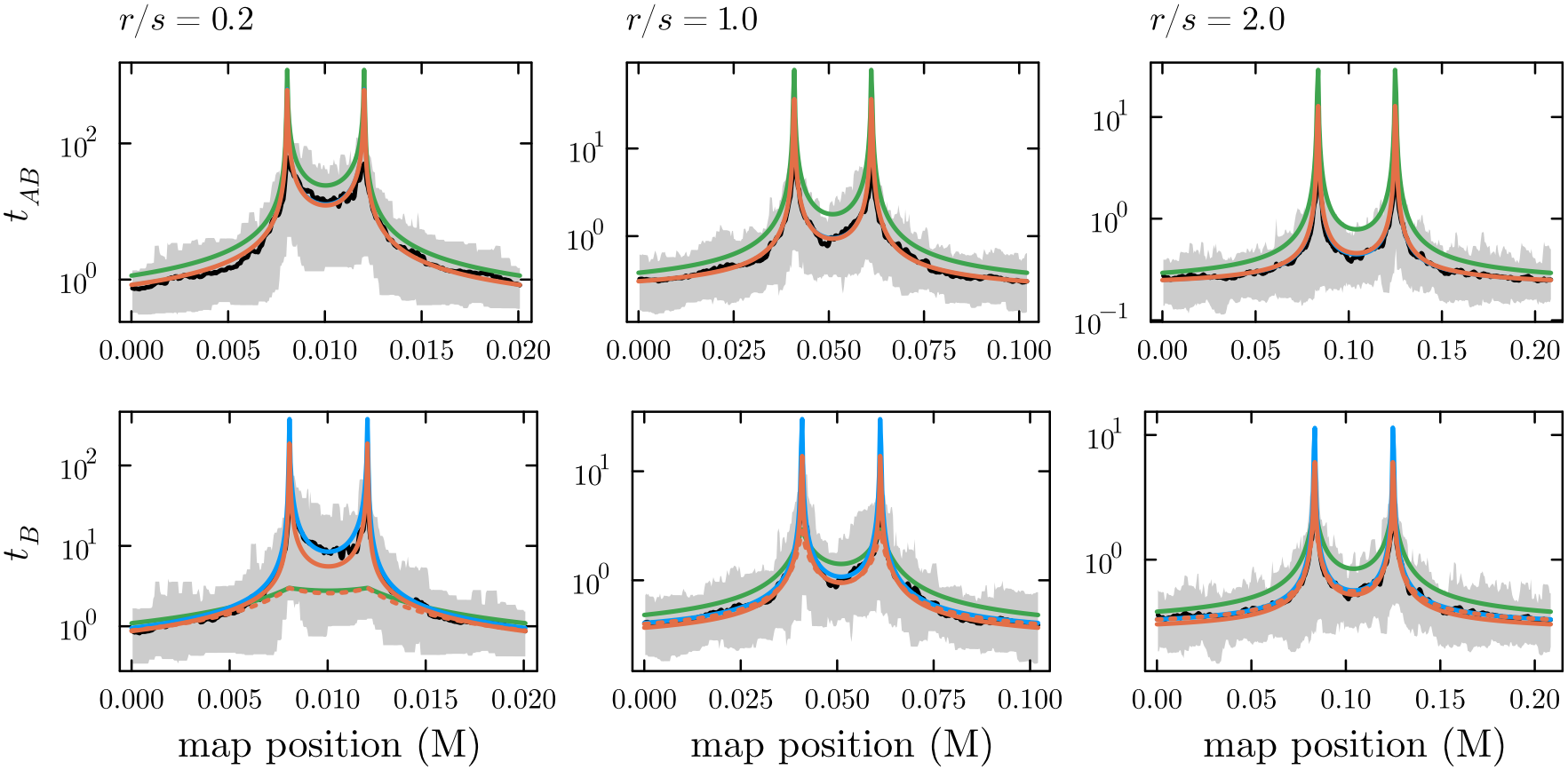
Between-(top row, *t*_*AB*_) and within-population (bottom row, *t*_*B*_) expected pairwise coalescence times for the model with two barrier loci. The black curve and shaded area show the mean and 95% density interval for the average pairwise coalescence time (in *N*_*B*_ generations) based on individual-based simulations (40 replicate simulations). The blue lines show theoretical predictions based on the structured coalescent model (section S1.2). The orange lines show predictions based on our new *m*_*e*_ approximation (eq. (7)), substituting *m*_*e*_ for *m* into the neutral mainland-island model coalescent prediction. The green lines show predictions based on Aeschbacher and Bürger (2014). Note that the structured coalescent prediction (blue) and the predictions based on our new *m*_*e*_ approximation are often almost indistinguishable (especially for *t*_*AB*_). The former relies on simulation-based estimates of barrier allele frequencies, whereas the latter co-estimates the allele frequencies and *m*_*e*_. For the within-population coalescence times, the solid orange lines show the *m*_*e*_-based prediction with the mixture model approximation (eq. (11)), while the dotted lines show predictions without the mixture model approximation (i.e. 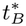 in eq. (11)). All results assume *N*_*B*_*s* = 10, *m/s* = 0.5 and *s* = 0.02.

As linkage become more tight, the region in between the two barrier loci becomes more and more differentiated relative to the genomic background, blurring the distinction between the barrier loci. This observation can be made somewhat more precise: denoting by *t*_*f*_ the between-population coalescence time in the flanking region a distance *r/*2 from the nearest selected locus and by *t*_*b*_ the between-population coalescence time at the midpoint in between the two barrier loci, we find that 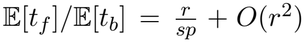 (using eqs. (4) and (5)). This lack of resolution will be exacerbated if the barrier loci have been subject to divergent selection for only a limited time (illustrated in fig. S10).

Again, we see clearly that a naive approach based on plugging *m*_*e*_ for *m* in the neutral prediction for the within-population coalescence time does not work when *r/s <* 1 (dashed lines in fig. 4). Nonetheless, the mixture approximation (eq. (11)) with the mixture proportion 2*h* predicted according to eq. (S23) appears to provide a fair approximation when linkage is tight.

### Polygenic barriers

We now proceed to consider polygenic barriers to gene flow, focusing on empirically plausible genomic architectures of divergent selection. A systematic exploration of the parameter space is challenging as it would involve varying the number of loci (*L*), their positions along the genome and selection coefficients (*x*_*i*_ and *s*_*i*_, for *i* = 1, … , *L*), the strength of drift (1*/N*_*B*_) and rate of migration (*m*). This does not appear feasible, so we zoom in on a couple of example simulations instead. Can we still predict adaptive divergence and neutral differentiation using *m*_*e*_ when divergent selection acts on many loci spread along the genome? What if selected loci are clustered in particular regions of the genome? How does the barrier effect change as the genetic architecture becomes more polygenic, i.e. when a given selective disadvantage of migrants is caused by more loci of smaller effect?

#### Predicting the genomic landscape of differentiation: a detailed simulation example

Consider the example displayed in fig. 5. Here we assume *L* = 200 divergently selected loci randomly scattered (uniformly) along a genome with three 100Mb chromosomes and a constant recombination rate of 1cM/Mb. This results in an average recombination rate 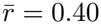 between pairs of selected loci, which should reflect a realistic genetic architecture: 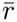 is estimated to be about 0.47 in humans and 0.30 in *Drosophila* (Veller et al., 2019, 2020). The selection coefficients are drawn from an exponential distribution of fitness effects with mean 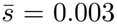, so that the total selection strength is about 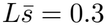 (i.e. when the island is fixed for the locally beneficial genotype, a migrant individual has a fitness reduction of about 26%). The island population size is chosen such that 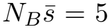 (i.e. *N*_*B*_ = 1667), and the mainland is assumed to be of the same size. We assume migration is unidirectional and fairly strong relative to selection, with 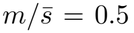 , corresponding to an expected 2.5 migrants per generation arriving on the island. Lastly, we assume a divergence time of *t*_*d*_ = 100.000 generations in the past, and an ancestral population of size *N*_*C*_ = *N*_*A*_ + *N*_*B*_.

**Figure 5.**
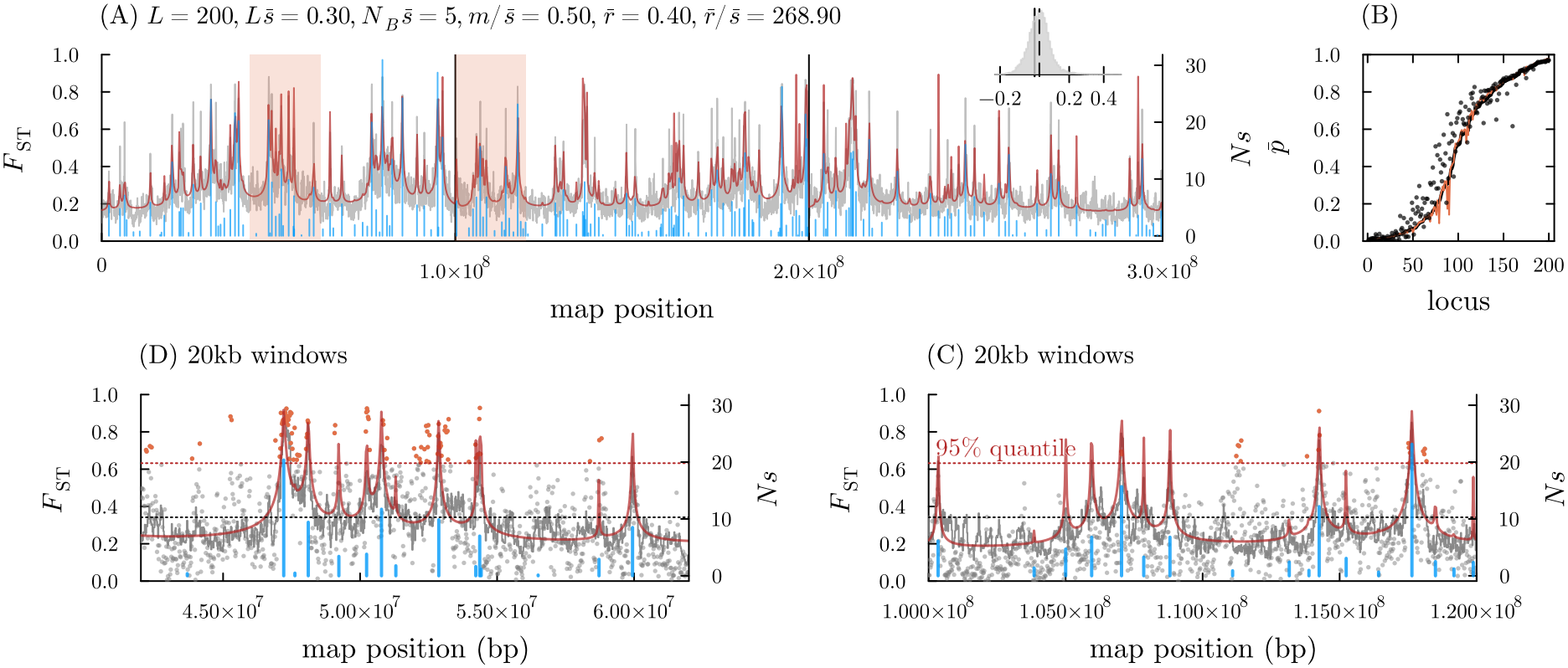
Genome-scale example with a polygenic barrier architecture. See main text for explanation of the parameterization. (A) Genomic landscape of differentiation (*F*_ST_ profile) along the genome. The gray line shows average *F*_ST_ along the genome estimated from five replicate simulations. The red line shows the *m*_*e*_-based prediction. Vertical blue lines mark divergently selected loci, with the height proportional to the selection coefficient (the *Ns* scale is shown on the right vertical axis). Black vertical lines delimit the three chromosomes. The highlighted areas are those shown in (C) and (D). The inset shows the distribution of the difference between the average estimated and predicted *F*_ST_ in 20kb windows, with the mean value marked by the dashed black line. (B) Allele frequencies at selected loci. The black line shows predicted allele frequencies from the *m*_*e*_ approach for all selected loci, ordered from low to high adaptive divergence. The red line shows predictions obtained using the theory of Zwaenepoel et al. (2024). Dots show the allele frequencies estimated from individual-based simulations, sorted by the predicted allele frequency. (C) Close-up view for the 42-62Mb region in (A), with dots showing average *F*_ST_ values estimated from neutral markers in 20kb windows from a sample of 20 haploid genomes in each population (assuming a neutral mutation rate of 10^*−*8^). The black horizontal line shows the genome-wide average *F*_ST_ for the 20kb windows. The red horizontal line shows the 95% quantile. Red dots show ‘outlier’ *F*_ST_ values, taking the 95% quantile as a threshold. (D) As in (C) but for the 100Mb to 120Mb region.

Our *m*_*e*_ approach predicts the observed *F*_ST_ profile (averaged across five replicate simulations, fig. 5A) and allele frequencies at selected loci remarkably well (fig. 5B). A slight upward bias is observed when comparing predicted *F*_ST_ values in 20kb windows against estimates from individual-based simulations (mean difference and SE of 0.025 *±* 0.0005, fig. 5A, inset). This bias seems to stem primarily from an underestimation of the within-island coalescence time close to barrier loci (fig. S11). If we compare *F*_ST_ estimates based on simulated SNPs in windows (instead of coalescent times observed in the recorded ARG) no such bias is apparent when compared against backward simulations based on the *m*_*e*_ approximation (fig. S12B,E) (In these backward simulations, SNP data is simulated in a window-by-window fashion using `msprime`, with the mean predicted *m*_*e*_ in the relevant window substituted for *m*). Furthermore, we find that not only the mean *F*_ST_ in windows, but also the variance thereof, matches very well with simulated data from the structured coalescent with mean *m*_*e*_ substituted for *m* (fig. S12C,F), although these estimates are rather noisy. Overall, these results suggest that the structured coalescent with a heterogeneous *m*_*e*_ along the genome provides a good model for the genomic landscape of differentiation when there is polygenic divergent selection.

Allele frequency divergence appears to be slightly underestimated by our theoretical predictions, especially for partially divergent loci (*p <* 0.8, say) (fig. 5B). The predicted allele frequencies are very similar to those obtained using the theory of Zwaenepoel et al. (2024). The gff for *selected* loci ranges from 0.08 to 0.54, with a mean value of 0.38. Effective gene flow at any selected locus is hence reduced by roughly two-to tenfold due to LD (i.e. selection on associated variants), depending on where in the genome it is located. The average *neutral* gff in 20kb windows ranges from 0.001 to 0.51, with a mean value of 0.29, which means that the time until a neutral lineage traces back to the mainland is increased by roughly two-to thousand-fold relative to the neutral expectation, again depending on genomic location.

This simulation further illustrates several important features. We see that an appreciable fraction of the divergently selected loci is not maintained divergently at equilibrium (i.e. is subject to swamping by gene flow), with the locally beneficial allele frequency below 0.025 for 10% of the loci, and below 0.1 for about 20% of the loci (fig. 5B). Obviously, such loci cannot be detected in an *F*_ST_-based genome scan even with perfect data (see examples in fig. 5C, D), showing how the genetic architecture of *reproductive isolation* (i.e. the *realized* barrier architecture, Zwaenepoel et al. (2024)) can be considerably different from the architecture of locally adaptive *traits*. This ‘hidden’ portion of trait architecture will depend strongly on the effective size of the island population and the strength of migration relative to selection.

Power to map barrier loci from outlier-scan approaches remains rather limited even when loci do maintain divergent alleles at equilibrium (and hence contribute to RI), because of variability in coalescence times, sampling variance of site-based statistics and the limited resolution due to the usage of window-based measures (Lohse, 2017). For instance, in a simulated data set with 20 haploid genomes sampled from each population and a neutral mutation rate of 10^*−*8^, none of the five barrier loci in the 105Mb to 110Mb region shown in fig. 5D are associated with outlier *F*_ST_ windows, despite equilibrium *F*_ST_ values that exceed the 95% percentile. Elsewhere in fig. 5C and D we see regions that are not quite close to any barrier locus that do show outlier *F*_ST_ windows.

Importantly, loci that maintain divergent alleles and contribute to the barrier to gene flow are not just those for which *s*_*i*_ *> m*, but include weakly selected loci that are protected from swamping due to the proximity of other, more strongly selected, barrier loci. As an example, consider the most weakly selected locus that maintains a 10% allele frequency difference between mainland and island (*p ≥* 0.1) in the simulation example of fig. 5. This locus has *m/s >* 2, but *g* = 0.08, leading to *m*_*e*_*/s* = 0.18 and an adaptive allele frequency of 0.21 on the island. This locus is rather tightly linked (*r ≈* 0.0005) to a more strongly selected locus (for which *m/s* = 0.36, and *m*_*e*_*/s* = 0.11), and lies within a 200kb region that contains three barrier loci in total (around the 54Mb point in fig. 5C). If this locus were unlinked to the rest of the barrier architecture, the associated gff would be 0.60, and the predicted allele frequency on the island would be 0.01. Arguably, this is an example of ‘divergence hitchhiking’ (Nosil, 2012), where differentiation at one locus is enabled by the close proximity of a barrier locus. One can use the ratio of the predicted barrier strength (*b* = *g*^*−*1^, Barton and Bengtsson (1986)) to the prediction *b*^*^ for the unlinked case (i.e. the barrier strength calculated for a locus with the same selection coefficient, but not linked to the other loci) as a quantitative measure of the strength of divergence hitchhiking relative to ‘genome hitchiking’. This is shown in fig. 6 (where the focal locus is marked by an arrow), highlighting the strong heterogeneity in hitchhiking effects along the genome. Figure 5C further shows that this 200kb stretch lies, in turn, within a larger *≈* 7Mb region of elevated differentiation that contains a number of strongly selected barrier loci.

**Figure 6.**
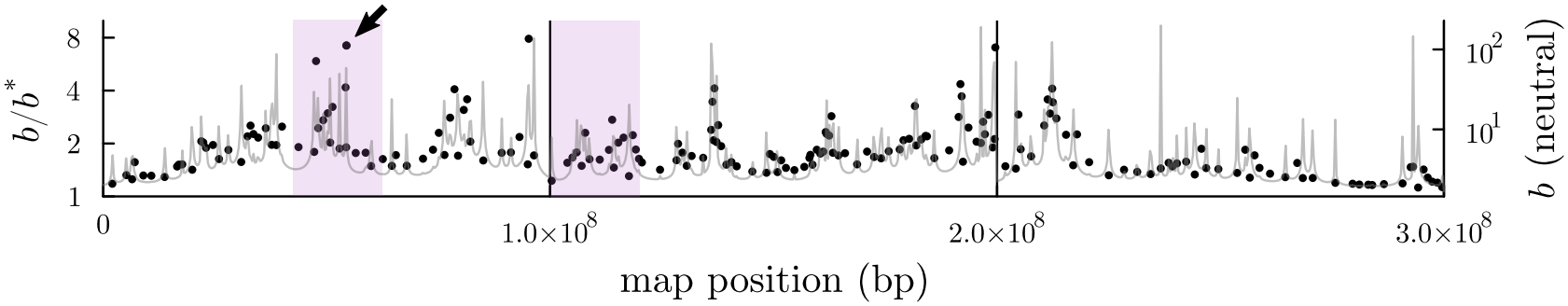
Quantifying divergence hitchhiking along the genome for the example barrier architecture of fig. 5. The black dots show the relative strength of divergence hitchhiking to genome hitchhiking at divergently selected loci as measured by *b/b*^*^. The gray line (right axis) shows the barrier strength at neutral sites along the genome. The shaded areas show the regions highlighted in fig. 5C and D. The focal locus discussed in the main text is marked by the black arrow.

#### Effects of linkage

The gff at any locus factors into a contribution from selected loci that are physically linked and a contribution from loci on different chromosomes (eq. (7)). Hence, given that eq. (7) appears to yield good predictions in simulation experiments with multiple chromosomes (fig. 5), we can focus on a single chromosome to examine in more detail the effects of linkage, absorbing selection at unlinked loci into a reduced migration rate parameter.

In fig. 7 we consider a genetic architecture of 100 selected loci scattered uniformly along a 100Mb region for three different genetic map lengths. Notably, predicted allele frequencies, coalescence times and *F*_ST_ remain remarkably accurate even for short map lengths. Again, we seem to slightly underpredict expected allele frequency divergence at barrier loci, especially for partially divergent loci (0.1 *< p <* 0.9, roughly). Predicted allele frequency distributions appear to match results from forward simulations fairly well (fig. S13). Interestingly, it appears that the lack of fit for some loci is not alleviated when fitting Wright’s distribution with two free parameters (*m*_*e*_ and *s*_*e*_) instead of one (*m*_*e*_), suggesting that marginal allele frequency distributions may no longer be well-described by the latter (in contrast to the two-locus case, fig. 3), and that a description based on effective parameters is no longer accurate. However, the allele frequency spectra estimates based on forward simulations are rather noisy, and this may also partly be responsible for the observed lack of fit.

**Figure 7.**
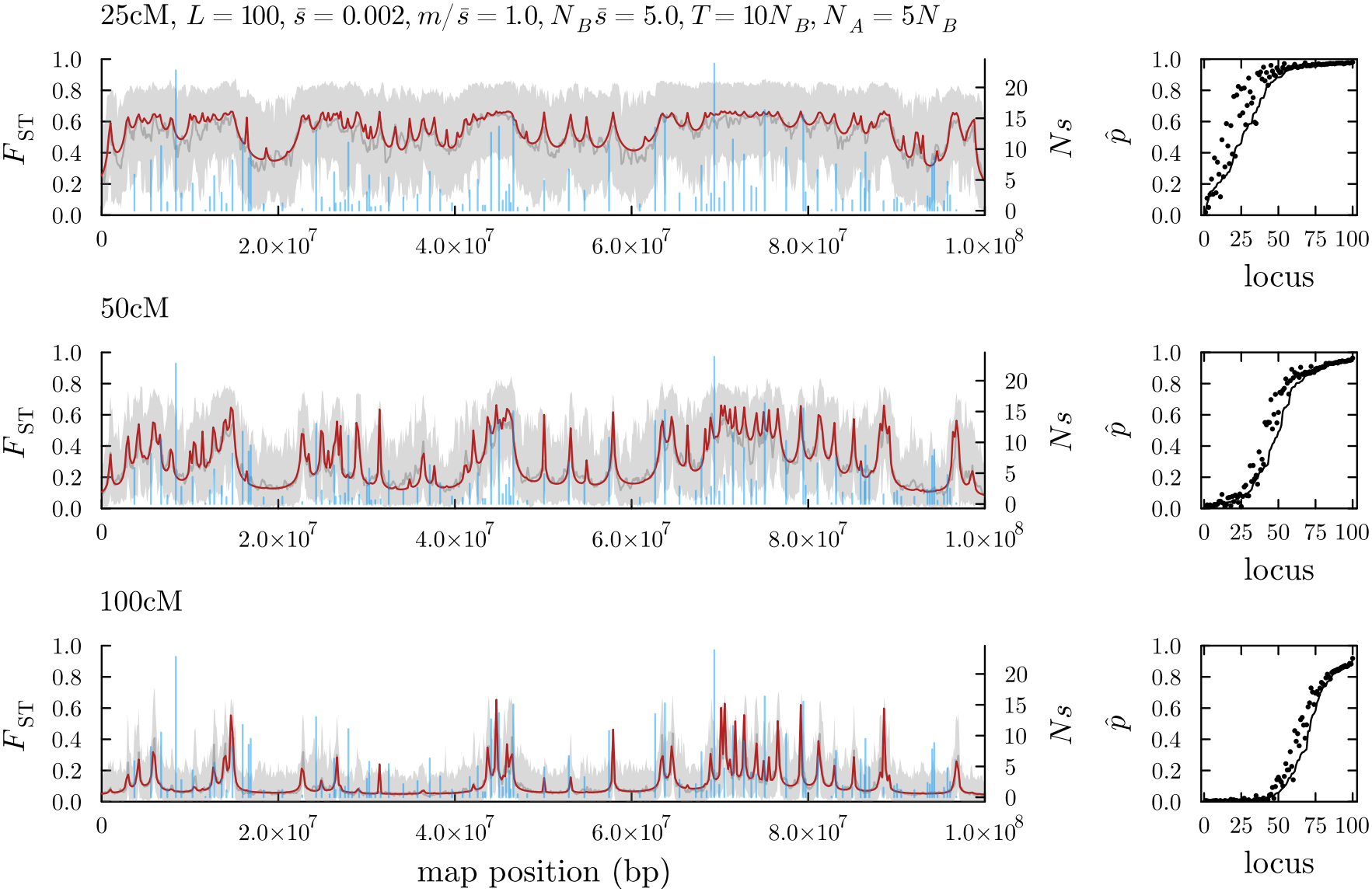
Effects of linkage on patterns of neutral differentiation along a single chromosome. From top to bottom results are shown for the same barrier architecture but with different associated map length (25, 50 and 100cM for the entire 100Mb region, respectively). The left plots show results from individual-based simulations (gray) and numerical predictions based on the *m*_*e*_ theory developed (red), with the vertical sticks showing the genomic positions of selected loci and their population-scaled selection coefficients (*Ns*, height of the sticks). The gray band shows the 95% density interval based on 10 replicate simulations, each with 10 replicate recapitations with 10 mainland sample lineages (see methods). The scatter plots on the right are as in fig. 5B). See also fig. S14.

As linkage becomes more tight, more and more barrier loci are able to maintain divergent alleles at equilibrium, increasing the neutral barrier to gene flow along the entire region, yielding a relative ‘flat’ *F*_ST_ profile. Note that the value at which *F*_ST_ levels out (here roughly 0.6) depends on the divergence time (which is identical to the timing of the onset of divergent selection in these simulations) and relative size of the mainland and island. When linkage is more loose (total map length is larger), fewer loci maintain divergent alleles, and the *F*_ST_ profile shows more clear ‘islands of divergence’: regions where multiple selected loci are rather tightly clustered and divergence hitchhiking plays an important role in shaping patterns of differentiation (see fig. S15 for a graph of *b/b*^*^ similar to fig. 6).

A uniform distribution of barrier loci along the genome may not be a realistic assumption. For instance, it is plausible that barrier loci are associated with coding regions, the density of which is typically rather heterogeneous along the genome. We refer to a barrier architecture where loci tend to be less evenly spaced on average than a uniform scattering as a ‘clustered’ architecture. While clustering does not affect the average recombination rate among pairs of loci, it can reduce the *harmonic mean* recombination rate among barrier loci strongly, which should result in a stronger barrier effect (Barton, 1983). In fig. S14, we show both simulations and predictions for a more clustered barrier architecture, and in fig. S15 we compare the extent of divergence hitchhiking between the uniform and clustered architecture. We find that clustering promotes adaptive divergence, i.e. more loci are maintained divergently between populations and are protected from swamping (figs. S14 and S16). This is because when barrier loci are clustered, divergence hitchhiking is more prominent: a weakly selected and swamping-prone locus is more likely to find itself in the vicinity of a strongly selected locus. For instance, for the 50cM map, the average value of *b/b*^*^ for the clustered architecture is almost 50 times that of the uniform one (fig. S15). However, the genome-wide average *neutral* gff does not change markedly when architectures are more clustered (fig. S15).

#### Effects of polygenicity

Now consider what happens as the barrier architecture on a single chromosome becomes more polygenic, i.e. when *L* increases while 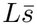 remains constant, so that the same fitness disadvantage in migrants is determined by more and more loci of smaller and smaller average effect 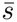. If we simultaneously hold *m* and *N*_*B*_ fixed, this implies that 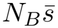 decreases while 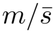 increases, so that we expect swamping to become more prevalent (fig. S17B). If, on the other hand, we hold 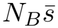 and 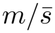 fixed (implying an increase in *N*_*B*_ with increasing polygenicity), drift and migration pressure *per locus* remain the same. In the latter scenario we see that the barrier to gene flow becomes stronger as the genetic architecture becomes more polygenic (fig. S17A). This appears to be driven by the decrease in the harmonic mean recombination rate among selected loci, which for a fixed map length is roughly proportional to 1*/* log *L* (Barton (1983)). If we scale the total map length by log *L* as we increase *L*, the distribution of allele frequency divergence across loci becomes largely independent of *L* (fig. S17C). As a consequence, the result of increasing polygenicity while maintaining a fixed effective population size is non-monotonic (fig. S17D): as the number of loci increases, the more tightly linked map first leads to a decrease in the average gff at selected loci (“the tightening of linkage outweighs the weakening of selection” – Barton (1983)). However as *L* becomes larger, and 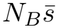 smaller, swamping eventually takes over and weakens the barrier to gene flow.

### Bidirectional migration

The assumption of unidirectional migration in the mainland-island model may appear as a significant limitation of the theory developed here. However, the fundamental assumption behind our approach is not unidirectional migration *per se*, but rather that migrants come from a pool with a known genetic composition. Consider for instance symmetric migration at rate *m*. The probability of a migrant or one of its descendants migrating back into the source population is *O*(*m*^2^). When migration is weak, this should be negligible, and one can think of the bidirectional migration case as a composition of two unidirectional mainland-island models, i.e. population *B* receives each generation an expected *mN*_*B*_ migrants that are population *A residents*, and *vice versa*. These two mainland-island models are however coupled: migration from *A* into *B* will affect the allele frequencies of divergently selected loci in the *B* resident gene pool, and hence will affect the genetic composition of the migrants *A* receives, in turn, from *B* (and *vice versa*). This suggests a fixed point iteration similar to the one for the mainland-island model, could be used to jointly predict allele frequencies in the two resident gene pools (see section S1.4).

While we leave a detailed study of the effects of bidirectional migration on patterns of differentiation and adaptive divergence to future work, we conducted a couple of simulations for the symmetric migration case to test our suggested approach. We find that we still obtain remarkably accurate predictions of both neutral *F*_ST_ and allele frequencies at selected loci as long as *m* is not too large (fig. 8). When *m* increases, the expected minor allele frequencies at selected loci increase to 0.5, and remain fairly accurately estimated by the *m*_*e*_-based approach. Note that these are *expected* allele frequency predictions: when *m > s*, the allele frequency distribution is often highly bimodal (‘U-shaped’), corresponding to a situation where the population spends extensive amounts of time being fixed for either one or the other allele. When *m* increases, neutral differentiation appears to be overpredicted. Note that our neutral predictions *do* take into account bidirectional migration, including *O*(*m*^2^) contributions to pairwise coalescence times (recall that we use phase-type theory for our calculations, with *m*_*e*_ substituted for *m*), so the source of this overprediction lies in the *m*_*e*_ approximation itself becoming poor, not the calculation of *F*_ST_.

**Figure 8.**
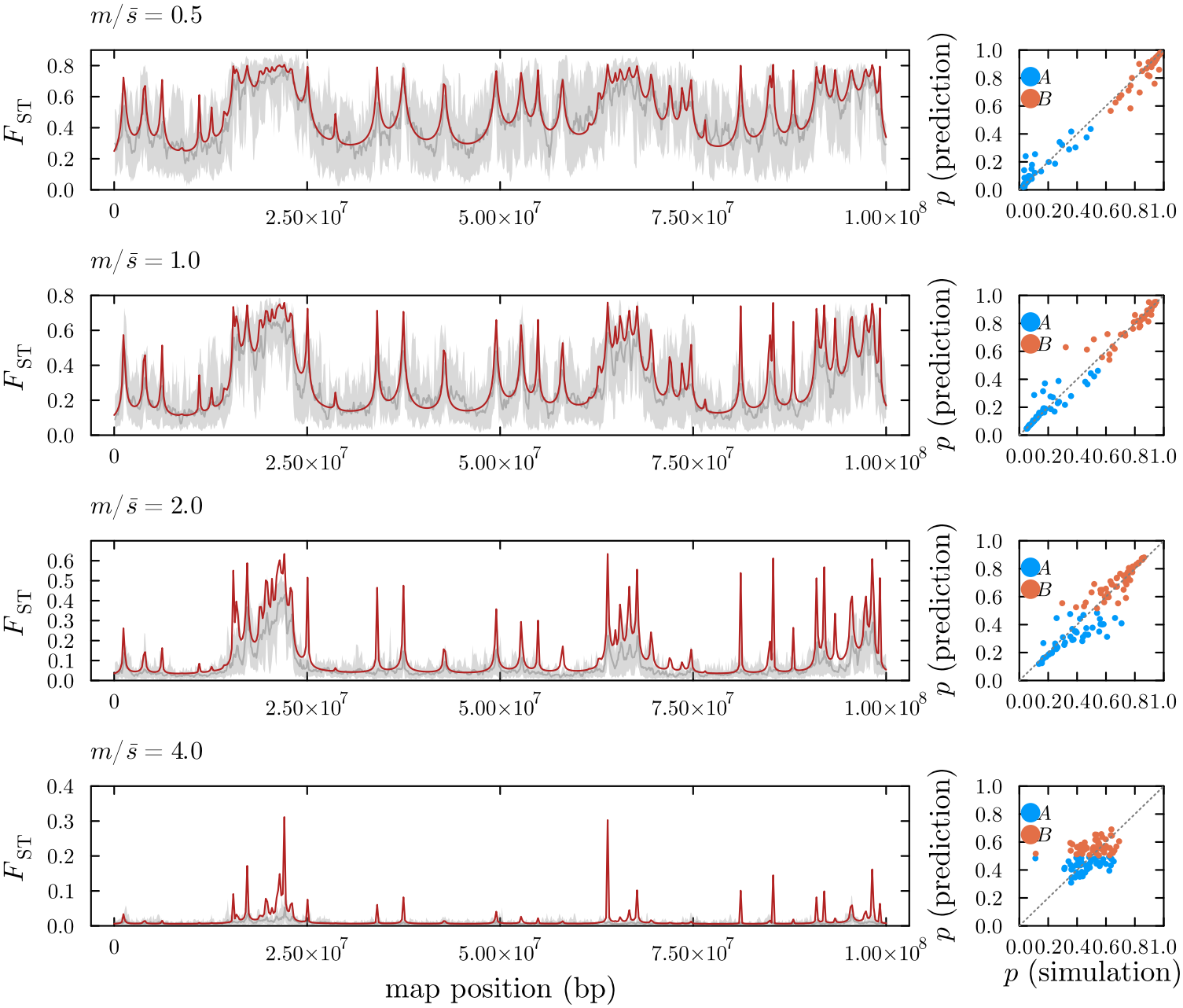
Predicted landscapes of differentiation (*F*_ST_, left column) and allele frequencies at selected loci (right column) for an IM model with symmetrical bidirectional migration. Each row shows predictions (red line) and estimates based on individual-based simulations (gray line, with 95% density interval) for the same genetic architecture, but with a different migration rate (shown as *m/s*). The scatterplots show predictions (*y*-axis) for expected allele frequencies at selected loci in both populations *A* and *B* against estimates from individual-based simulations (*x*-axis). We assume a 50cM, 100Mb genetic map with *L* = 50, 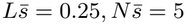 , *N*_*A*_ = *N*_*B*_ = *N* = 1000, *t*_*d*_ = *t*_*c*_ = 10*N* . Results are based on 10 replicate simulations.

## Discussion

In this paper, we present a new approach for calculating effective migration rates along the genome for the mainland-island model with polygenic barriers to gene flow. We show how to use this *m*_*e*_ for predicting adaptive divergence at barrier loci and expected coalescence times at linked neutral loci in IM-like demographic models. Our *m*_*e*_ improves considerably on previous approximations: like Aeschbacher and Bürger (2014) (see also Aeschbacher et al. (2017)), we obtain accurate predictions for neutral *m*_*e*_ even when barrier loci are tightly linked; however, unlike the latter authors, we do not assume *m/s* ≪ 1 (i.e. we do not assume complete divergence at selected loci), and as in Zwaenepoel et al. (2024) and Sachdeva (2022), we jointly predict *m*_*e*_ and allele frequency divergence between mainland and island at equilibrium. The latter authors, however, assumed loose linkage. Using this novel *m*_*e*_ approximation, we are able to predict the genomic landscape of differentiation for a given genetic architecture of divergent selection with unprecedented accuracy across almost the entire biologically relevant parameter range.

### Effectiveness of effective migration rates

Expected allele frequencies at barrier loci are predicted accurately by our *m*_*e*_-based approach, even when linkage among barrier loci becomes tight. However, in the latter case, the allele frequency *distribution* for the two-locus model suggests that a parameterization with an effective selection coefficient is often more adequate. Whether more adequate approximations can be worked out that make use of both an effective selection coefficient and migration rate remains an open question.

Our *m*_*e*_-based approximation provides excellent predictions for the *between*-population coalescence time at a neutral locus associated with a barrier, even when linkage is tight and migration strong. The situation for *within*-population coalescence times is somewhat different. When linkage is sufficiently loose, the latter remain well-described by the neutral structured coalescent with *m*_*e*_ substituted for *m*. However, for neutral loci tightly linked to a barrier, the within-island coalescence time is underestimated by the naive approach of substituting *m*_*e*_ for *m* in the neutral prediction, especially when there is partial divergence (*m* is not much smaller than *s*). This is a result of (genetic) class structure in the island population (fig. 1A). In models with one or two barrier loci, this class structure can be explicitly accounted for in a (doubly) structured coalescent model (Nordborg, 1997) (see sections S1.1 and S1.2), but this does not extend easily to a polygenic model. We found that a mixture approximation is able to correct remarkably well for the error in predicted within-population coalescence times, both for models with one or two barrier loci and the polygenic case. While promising, predicting the mixture proportion remains rather *ad hoc* (see section S1.5) and should be a target of further investigation.

Importantly, making use of recent developments in phase-type theory (Hobolth et al., 2024; Sendrowski and Hobolth, 2026), we show how our *m*_*e*_ approximation can be used to obtain accurate predictions for rather general demographic models, beyond the rather limiting mainlandisland model on which the theory is based. Notably, we find that our modeling approach extends rather well to the case with bidirectional migration, although we have not studied this in a very systematic way. While we have only exclusively focused on pairwise coalescence times, the approach based on phase-type theory in principle allows us to obtain coalescence time predictions for arbitrary sampling configurations. However, one would again have to deal with the problem of class structure when linkage is tight. A more general solution for this issue (i.e. beyond our mixture approximation) would therefore be welcome.

### The genetic architecture of RI with polygenic divergent selection

Our results provide insights into how the genetic architecture of divergent selection, genetic drift and migration shape the barrier to gene flow along the genome. Specifically, we are able to predict when alleles that are under divergent selection pressures can contribute to RI, and when they are likely to be swamped by gene flow. We show through examples how the ability of a locus to contribute to RI at equilibrium can depend strongly on genomic context, and suggest a measure to quantify the strength of such ‘divergence hitchhiking’ effects (Nosil, 2012). Notably, even when selected loci are uniformly scattered along the genome, appreciable heterogeneity in the extent of divergence hitchhiking along the genome is observed (fig. 6, fig. S15). When barrier loci are clustered along the genome, the extent of divergence hitchhiking increases, and more weakly selected loci can contribute to RI at equilibrium, while the genome-wide neutral barrier to gene flow is not much affected. The effects of polygenicity *per se* were also briefly considered. While barrier strength is largely independent of *L* in the unlinked setting as long as 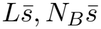 and 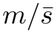 are held constant (Sachdeva, 2022; Zwaenepoel et al., 2024), when the map length is held fixed, increasing *L* necessarily leads to a decreased harmonic mean recombination rate and increased barrier strength. However, when *N*_*B*_ is held fixed with increasing polygenicity, we see a more complicated picture: as *L* increases (keeping 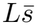 constant), tighter coupling first yields a stronger barrier to gene flow, until 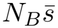 becomes too low to prevent swamping by gene flow.

### Implications for genome scan approaches

Empiricists often calculate site-based measures of differentiation such as *F*_ST_ in windows along the genome in order to map regions involved in local adaptation and species barriers (Ravinet et al., 2017). Our approach for jointly predicting expected coalescence times and allele frequencies at barrier loci enables predicting the expected outcome of such a genomic scan for a *given* genetic architecture and demographic model from first principles. While our *m*_*e*_ approach enables accurate prediction of site-based statistics and hence *expected values* of window-based statistics, it does not provide a means to calculate the *correlation* between site-based statistics along the genome, and hence does not enable calculation of the *variance* of window-based statistics. Nevertheless, backward simulations in windows along the genome with the mean *m*_*e*_ substituted for *m* appear to yield a similar variance for the mean *F*_ST_ in windows (fig. S12). In principle, then, our theory could be used to answer questions related to the power of genome scan approaches, e.g. “For a given architecture of divergent selection, effective population size, migration rate and sample size, what is the power of an outlier-based *F*_ST_ scan? What is the false positive rate?” However, this is perhaps not that useful, as such questions rely on postulating a *particular* genetic architecture, while the space of possible architectures is vast. In general, the inability to specify a pertinent null model when divergent selection is polygenic makes it difficult to put outlier scan approaches on a more statistically controlled footing.

### Implications for model-based inference of barrier architecture

The finding that the distribution of pairwise coalescence times is usually well-approximated by a neutral model with an effective migration rate suggests that site-based measures of population differentiation do not provide much information about the genetic architecture (i.e. the number of selected loci, their genomic locations and fitness effects) *beyond m*_*e*_. This in turn suggests that when we aim to learn about the genetic architecture of reproductive isolation from neutral SNP data, the relevant question is “which genetic architectures are compatible with a particular *m*_*e*_ profile and *N*_*e*_?”^2^.

An approach like the one of Laetsch et al. (2022) therefore appears sensible: first fit a neutral IM model in windows along the genome, with *m*_*e*_ and *N*_*e*_ parameters that are allowed to vary across windows, and afterwards fit a model of barrier architecture (they used the model of Aeschbacher et al. (2017)), treating the estimated *m*_*e*_-profile as data. However, such a two-step approach ignores uncertainty in the *m*_*e*_ and *N*_*e*_ estimates, and does not harness the fact that *m*_*e*_ is correlated along the genome, which entails that the value of *m*_*e*_ in nearby genomic regions provides information about plausible *m*_*e*_ values at any particular position. Overall, there is a rather complicated dependence: *m*_*e*_ at any given genomic position depends on allele frequencies at all barrier loci, which themselves depend on *m*_*e*_, *N*_*e*_, *s* and the genomic locations of those barrier loci. Inference of barrier architecture from neutral polymorphism data may therefore benefit from a joint inference approach, in which prior constraints on e.g. distributions of fitness effects, migration rates, genomic locations of barrier loci and *N*_*e*_ mutually constrain the space of plausible genetic architectures. The *m*_*e*_ theory presented here should be useful to develop such an approach, not unlike inference machinery developed for linked selection within a single population (Elyashiv et al., 2016; Murphy et al., 2022; Buffalo and Kern, 2024). Clearly, such an inference approach would have to account for other forms of linked selection if it is to be useful for mapping barriers to gene flow (Cruickshank and Hahn, 2014).

While the above considerations apply to site-based measures, it is currently unclear whether haplotype-based information would significantly alter our power to map the genetic architecture of reproductive isolation. In general, approaches modeling selection using effective parameters, whether *m*_*e*_ or *N*_*e*_, remain stuck in a site-by-site perspective of genetic variation along the genome, providing approximations for *marginal* coalescence time or allele frequency distributions at any given position. Whether approaches based on effective parameters could be extended to derive joint predictions for multiple sites remains an open question.

### Limitations

Other processes than divergent selection affect the genomic landscape of differentiation (Cruickshank and Hahn, 2014; Schield et al., 2025). Background selection (BGS), in particular, is ubiquitous and heterogeneous along the genome, and has been subject to considerable recent interest in the context of genomic landscapes of differentiation (Rodrigues et al., 2024; Schield et al., 2025; Gabrielli et al., 2026). BGS can often be reasonably well-modelled by a reduction in *N*_*e*_ that varies along the genome which can be predicted for a given genetic architecture of purifying selection (Cvijović et al., 2018; Santiago and Caballero, 2016; Buffalo and Kern, 2024). How BGS affects genetic variation and differentiation across populations with ongoing gene flow remains somewhat poorly understood. Both simulations and theoretical work suggest that, in the absence of divergent selection, the effects of BGS depend on the relative strength of migration and purifying selection (Matthey-Doret and Whitlock, 2019; Hasan and Whitlock, 2024). When migration is weak relative to purifying selection, classical BGS predictions of *N*_*e*_ should work well to predict the effects of BGS on genetic differentiation, while for strong migration BGS will often not have much of an effect on measures of differentiation such as *F*_ST_ (Hasan and Whitlock, 2024). However, the case with both BGS and divergent selection has, to our knowledge, not yet been considered. Whether and in what regimes BGS can be largely captured by a simple reduction in *N*_*e*_, which would then feed back into allele frequency predictions and *m*_*e*_, remains to be studied in detail.

Selective sweeps, possibly at barrier loci, constitute another form of linked selection that affects the landscape of genomic differentiation (Rodrigues et al., 2024; Schield et al., 2025). While our modeling approach accounts for the effects of selection on linked neutral variation at *equilibrium*, we have completely ignored the *transient* effects of positive selection (divergent or otherwise). Relatedly, we have assumed a scenario of secondary contact, where two populations that maintain alternative alleles at a set of divergently selected loci come into contact and evolve towards multilocus migration-selection equilibrium. We have completely ignored the process by which the two populations become divergent at these loci in the first place. Our model should readily apply in the case of a rapid polygenic response from standing variation in allopatry. However, when there is divergence with gene flow, the timescale at which multilocus migration-selection balance is attained becomes an important factor in determining when our modeling approach is reasonable (Yeaman and Whitlock, 2011).

For simplicity, we have only considered haploids, but extending the theory to deal with diploids and dominance (as in Zwaenepoel et al. (2024)) should not present too much difficulty. Importantly, we have assumed a very simple genotype-to-fitness map, involving directional selection on an additive trait. Loci with epistatic fitness effects (e.g. Dobzhansky-Müller incompatibilities) are of course expected to play an important role in the evolution of RI (Schumer et al., 2018) and may be associated, causally or not, with divergent adaptation (Bierne et al., 2011). Although one could extend the predictions for the two-barrier case to account for epistatic interactions (as Pontz and Bürger (2022) did for invasion probabilities in a similar two-locus system), the expressions for *m*_*e*_ would then depend not only on *p*_1_ and *p*_2_, but also on *D* (i.e. LD between the barrier loci). No good approximations exist for the latter outside of the deterministic case, so that our approach for co-predicting allele frequencies and *m*_*e*_ using the diffusion approximation and fixed point iteration would no longer be feasible. Accounting for the effects of multi-locus interactions in *m*_*e*_-based approximations hence remains an open problem. Additionally, divergent selection may involve stabilizing selection to alternative phenotypic optima, potentially leading to a rather different polygenic selection response compared to the one assumed here (Yeaman, 2015; Hayward and Sella, 2022). Understanding the genetic architecture of RI for more realistic models of divergent adaptation, where fitness is determined by stabilizing selection on multiple traits, remains a key challenge (Li et al., 2026).

## Data availability

The author affirms that all data necessary for confirming the conclusions of the article are present within the article, figures and supplementary material. Code for conducting the simulations and generating the figures in the main text and supplementary material is available at https://codeberg.org/arzwa/barriers-paper.

## Acknowledgments

Many thanks to Christelle Fraïsse, Himani Sachdeva, Curro Campuzano Jimenez and Hannes Svardal for feedback on draft versions of the present article. Arthur Zwaenepoel acknowledges funding from the Research Foundation – Flanders (FWO, Junior Postdoctoral Fellowship 1272625N).

## Footnotes

1 To get an idea of the physical distance associated with e.g. *r/s* = 0.1: if we assume *N*_*e*_ = 10^6^ and a recombination rate of 1cM/Mb, this would correspond to 100bp between the barrier loci for *N*_*e*_*s* = 10, 500bp for *N*_*e*_*s* = 50 and 1kb for *N*_*e*_*s* = 100. For *N*_*e*_ = 10^5^, this is 1kb, 5kb and 10kb, *etc*.

2 The error in the naive *m*_*e*_ prediction for within-population coalescence times (which we adjust using the mixture approximation) does provide some information about the rate at which migrant ancestry is purged (see section S1.5), which in turn is informative about some aspects of the genetic architecture (notably the average recombination rate between selected loci).

## Notes

### Competing Interest Statement

The authors have declared no competing interest.

### Summary of Updates

Minor revisions to enhance figure readability. A typo in the caption of figure 1. An additional remark in discussion on modeling epistasis, with a reference to Pontz & Burger (2022).

## References

S. Aeschbacher and R. Bürger. The effect of linkage on establishment and survival of locally beneficial mutations. Genetics, 197(1):317–336, 2014.

S. Aeschbacher, J. P. Selby, J. H. Willis, and G. Coop. Population-genomic inference of the strength and timing of selection against gene flow. Proceedings of the National Academy of Sciences, 114(27):7061–7066, 2017.

N. H. Barton. Multilocus clines. Evolution, pages 454–471, 1983.

N. H. Barton and B. O. Bengtsson. The barrier to genetic exchange between hybridising populations. Heredity, 57(3):357–376, 1986.

F. Baumdicker, G. Bisschop, D. Goldstein, G. Gower, A. P. Ragsdale, G. Tsambos, S. Zhu, B. Eldon, E. C. Ellerman, J. G. Galloway, et al. Efficient ancestry and mutation simulation with msprime 1.0. Genetics, 220(3):iyab229, 2022.

B. Bengtsson. The flow of genes through a genetic barrier. Evolution: essays in honour of John Maynard Smith, 1:31–42, 1985.

N. Bierne, J. Welch, E. Loire, F. Bonhomme, and P. David. The coupling hypothesis: why genome scans may fail to map local adaptation genes. Molecular ecology, 20(10):2044–2072, 2011.

V. Buffalo and A. D. Kern. A quantitative genetic model of background selection in humans. Plos Genetics, 20(3):e1011144, 2024.

E. Burban, M. I. Tenaillon, and S. Glémin. Ridge, a tool tailored to detect gene flow barriers across species pairs. Molecular Ecology Resources, 24(4):e13944, 2024.

R. Bürger and A. Akerman. The effects of linkage and gene flow on local adaptation: a two-locus continent–island model. Theoretical population biology, 80(4):272–288, 2011.

T. E. Cruickshank and M. W. Hahn. Reanalysis suggests that genomic islands of speciation are due to reduced diversity, not reduced gene flow. Molecular ecology, 23(13):3133–3157, 2014.

I. Cvijović, B. H. Good, and M. M. Desai. The effect of strong purifying selection on genetic diversity. Genetics, 209(4):1235–1278, 2018.

E. Elyashiv, S. Sattath, T. T. Hu, A. Strutsovsky, G. McVicker, P. Andolfatto, G. Coop, and G. Sella. A genomic map of the effects of linked selection in drosophila. PLoS genetics, 12 (8):e1006130, 2016.

C. Fraïsse, I. Popovic, C. Mazoyer, B. Spataro, S. Delmotte, J. Romiguier, E. Loire, A. Simon, N. Galtier, L. Duret, et al. Dils: Demographic inferences with linked selection by using abc. Molecular Ecology Resources, 21(8):2629–2644, 2021.

M. Gabrielli, T. Leroy, C. Roux, B. Milá, C. Thébaud, and B. Nabholz. Accounting for recombination rate variation improves inference of barrier loci and reveals the role of both natural and sexual selection in an incipient bird radiation. Evolution, page qpag121. 07 2026. ISSN 0014-3820. doi: 10.1093/evolut/qpag121. URL https://doi.org/10.1093/evolut/qpag121.

B. C. Haller, J. Galloway, J. Kelleher, P. W. Messer, and P. L. Ralph. Tree-sequence recording in slim opens new horizons for forward-time simulation of whole genomes. Molecular ecology resources, 19(2):552–566, 2019.

A. Hasan and M. C. Whitlock. Fst and genetic diversity in an island model with background selection. PLoS genetics, 20(12):e1011225, 2024.

L. K. Hayward and G. Sella. Polygenic adaptation after a sudden change in environment. Elife, 11:e66697, 2022.

J. Hey and R. Nielsen. Multilocus methods for estimating population sizes, migration rates and divergence time, with applications to the divergence of drosophila pseudoobscura and d. persimilis. Genetics, 167(2):747–760, 2004.

A. Hobolth, I. Rivas-González, M. Bladt, and A. Futschik. Phase-type distributions in mathematical population genetics: An emerging framework. Theoretical population biology, 157: 14–32, 2024.

R. R. Hudson and N. L. Kaplan. Deleterious background selection with recombination. Genetics, 141(4):1605–1617, 1995.

I. Juric, S. Aeschbacher, and G. Coop. The strength of selection against neanderthal introgression. PLoS genetics, 12(11):e1006340, 2016.

J. Kelleher, K. R. Thornton, J. Ashander, and P. L. Ralph. Efficient pedigree recording for fast population genetics simulation. PLoS computational biology, 14(11):e1006581, 2018.

Y. Kobayashi, P. Hammerstein, and A. Telschow. The neutral effective migration rate in a mainland-island context. Theoretical Population Biology, 74(1):84–92, 2008.

D. R. Laetsch, G. Bisschop, S. H. Martin, S. Aeschbacher, D. Setter, and K. Lohse. Demo-graphically explicit scans for barriers to gene flow using gimble. bioRxiv, pages 2022–10, 2022.

J. Li, J. Hermisson, and H. Sachdeva. Effect of population structure and stabilizing selection on quantitative genetic variation. bioRxiv, pages 2026–03, 2026.

K. Lohse. Come on feel the noise-from metaphors to null models. J. Evol. Biol, 30:1506–1508, 2017.

R. Matthey-Doret and M. C. Whitlock. Background selection and fst: consequences for detecting local adaptation. Molecular ecology, 28(17):3902–3914, 2019.

D. A. Murphy, E. Elyashiv, G. Amster, and G. Sella. Broad-scale variation in human genetic diversity levels is predicted by purifying selection on coding and non-coding elements. Elife, 12:e76065, 2022.

R. Nielsen, A. H. Vaughn, and Y. Deng. Inference and applications of ancestral recombination graphs. Nature Reviews Genetics, 26(1):47–58, 2025.

M. Nordborg. Structured coalescent processes on different time scales. Genetics, 146(4):1501–1514, 1997.

M. Nordborg, B. Charlesworth, and D. Charlesworth. The effect of recombination on background selection. Genetics Research, 67(2):159–174, 1996.

P. Nosil. Ecological speciation. Oxford University Press, 2012.

D. Petry. The effect on neutral gene flow of selection at a linked locus. Theoretical population biology, 23(3):300–313, 1983.

M. Pontz and R. Bürger. The effects of epistasis and linkage on the invasion of locally beneficial mutations and the evolution of genomic islands. Theoretical Population Biology, 144:49–69, 2022.

M. Ravinet, R. Faria, R. Butlin, J. Galindo, N. Bierne, M. Rafajlović, M. Noor, B. Mehlig, and A. Westram. Interpreting the genomic landscape of speciation: a road map for finding barriers to gene flow. Journal of evolutionary biology, 30(8):1450–1477, 2017.

M. F. Rodrigues, A. D. Kern, and P. L. Ralph. Shared evolutionary processes shape landscapes of genomic variation in the great apes. Genetics, 226(4):iyae006, 2024.

F. Rousset. Genetic structure and selection in subdivided populations, volume 40. Princeton University Press, 2004.

H. Sachdeva. Reproductive isolation via polygenic local adaptation in sub-divided populations: Effect of linkage disequilibria and drift. PLoS genetics, 18(9):e1010297, 2022.

E. Santiago and A. Caballero. Joint prediction of the effective population size and the rate of fixation of deleterious mutations. Genetics, 204(3):1267–1279, 2016.

D. R. Schield, J. K. Carter, M. G. Alderman, K. Farleigh, D. K. Highland, and R. J. Safran. Recombination rate and recurrent linked selection shape correlated genomic landscapes across a continuum of divergence in swallows. Molecular Ecology, 34(22):e70074, 2025.

M. Schumer, C. Xu, D. L. Powell, A. Durvasula, L. Skov, C. Holland, J. C. Blazier, S. Sankararaman, P. Andolfatto, G. G. Rosenthal, et al. Natural selection interacts with recombination to shape the evolution of hybrid genomes. Science, 360(6389):656–660, 2018.

O. Seehausen, R. K. Butlin, I. Keller, C. E. Wagner, J. W. Boughman, P. A. Hohenlohe, C. L. Peichel, G.-P. Saetre, C. Bank, Å. Brännström, et al. Genomics and the origin of species. Nature Reviews Genetics, 15(3):176–192, 2014.

J. Sendrowski and A. Hobolth. Phasegen: exact solutions for time-inhomogeneous multivariate coalescent distributions under diverse demographies. Genetics, 232(1):iyaf135, 2026.

M. Slatkin. Inbreeding coefficients and coalescence times. Genetics Research, 58(2):167–175, 1991.

S. Stankowski, M. A. Chase, H. McIntosh, and M. A. Streisfeld. Integrating top-down and bottom-up approaches to understand the genetic architecture of speciation across a monkeyflower hybrid zone. Molecular Ecology, 32(8):2041–2054, 2023.

C. Veller, N. Kleckner, and M. A. Nowak. A rigorous measure of genome-wide genetic shuffling that takes into account crossover positions and mendel’s second law. Proceedings of the National Academy of Sciences, 116(5):1659–1668, 2019.

C. Veller, N. B. Edelman, P. Muralidhar, and M. A. Nowak. Variation in genetic relatedness is determined by the aggregate recombination process. Genetics, 216(4):985–994, 2020.

C. Veller, N. B. Edelman, P. Muralidhar, and M. A. Nowak. Recombination and selection against introgressed dna. Evolution, 77(4):1131–1144, 2023.

A. M. Westram, S. Stankowski, P. Surendranadh, and N. Barton. What is reproductive isolation? Journal of evolutionary biology, 35(9):1143–1164, 2022.

J. B. Wolf and H. Ellegren. Making sense of genomic islands of differentiation in light of speciation. Nature Reviews Genetics, 18(2):87–100, 2017.

S. Wright. The distribution of gene frequencies in populations. Proceedings of the National Academy of Sciences, 23(6):307–320, 1937.

S. Yeaman. Local adaptation by alleles of small effect. The American Naturalist, 186(S1): S74–S89, 2015.

S. Yeaman and M. C. Whitlock. The genetic architecture of adaptation under migration– selection balance. Evolution, 65(7):1897–1911, 2011.

A. Zwaenepoel, H. Sachdeva, and C. Fraïsse. The genetic architecture of polygenic local adaptation and its role in shaping barriers to gene flow. Genetics, page iyae140, 2024.

