## Supplementary information for "Predicting barrier architecture and the genomic landscape of differentiation under polygenic divergent selection"

Arthur Zwaenepoel*[1]

[1]Department of Biology, University of Antwerp, 2020 Antwerp, Belgium

**S2 Supplementary figures** **12**

### S1 Supplementary information

#### S1.1 Backward-time analysis for a single barrier locus

Let $p$ be the frequency of the `0` allele (locally beneficial on the island) and $q = 1-p$ the frequency of the `1` allele. Using the Markov property, the expected time $\mathbb{E}[t_i]$ until a neutral lineage in genetic background $i$ is absorbed into the mainland can be found using the following 'first-step analysis' (see e.g. Nordborg (1997))

$$\begin{aligned}\mathbb{E}[t_0] &= \frac{1}{rq} + \mathbb{E}[t_1] \\ \mathbb{E}[t_1] &= \frac{1}{rp + m/q} + \frac{rp}{rp+m/q}\mathbb{E}[t_0]\end{aligned} \tag{S1}$$

Note that since the mainland is fixed for the `1` allele, only neutral lineages that are on the `1` background can move into the mainland. Solving this system yields the following expression for the expected absorption time for a lineage at a neutral locus linked to a migration-selection polymorphism

$$\mathbb{E}[t] = p\mathbb{E}[t_0] + q\mathbb{E}[t_1] = \frac{1}{r}\frac{p}{q} + \frac{1}{m} \tag{S2}$$

We can hence define an $m_e$ at the neutral locus as

$$m_e = \frac{1}{\mathbb{E}[t]} = m\frac{rq}{mp+rq} \tag{S3}$$

*

Assuming deterministic migration-selection balance ($q = m/s$), the gff can be expressed as

$$g := \frac{m_e}{m} = \frac{r}{r + sp} = \frac{r}{r + s - m}$$

For weak migration ($m \ll s, r$), this is approximately $r/(r + s)$, a result first obtained by other means in Petry (1983).

**Coalescence times** Let $A$ denote the mainland population of size $N_A$, and $B$ the island population of size $N_B$. The expected coalescence time within the mainland is $\mathbb{E}[t_A] = N_A$. The expected cross-population coalescence time is $\mathbb{E}[t_{AB}] = \mathbb{E}[t] + N_A$, with $\mathbb{E}[t]$ given by eq. (S2). To calculate $\mathbb{E}[t_B]$, the expected coalescence time within the island, let $P_{ij}$ be the probability that a neutral lineage currently in genetic background $i$ was in background $j$ one generation before. We fix the following ordering of genetic backgrounds (allelic states at the selected locus in a particular population):

1. `0` (locally beneficial) on the island
2. `1` (locally deleterious) on the island
3. `1` on the mainland.

Recall that we assume the mainland is fixed for the `1` allele, hence there is no `0` background on the mainland. The backward transition matrix is then

$$P = \begin{pmatrix} 1 - rq & rq & 0 \\ rp & 1 - rp - \frac{m}{q} & \frac{m}{q} \\ 0 & 0 & 1 \end{pmatrix} \tag{S4}$$

Given $P$, one can calculate the expected within-island coalescence time $\mathbb{E}[t_B]$ (see eq. (S9) below). This yields a complicated expression that does not bring much additional insight. We can, however, look at some interesting limit cases. As $r/s$ becomes large (i.e. the neutral allele is not tightly linked to the barrier), we find, as expected, that the within-island coalescence time is just the neutral expectation, i.e.

$$\mathbb{E}[t_B] = \frac{N_B\,(3 + 2mN_A)}{1 + 2mN_B} + \mathcal{O}(s/r), \quad s/r \to 0$$

On the other hand, when linkage becomes tight, we find that

$$\begin{aligned} \mathbb{E}[t_B] &= \frac{2(s - m)}{N_B r s} + \mathcal{O}(1), \quad r/s \to 0 \\ &= \frac{2p}{N_B r} + \mathcal{O}(1), \quad r/s \to 0 \end{aligned}$$

Hence, if we keep $q = 1 - p = m/s$ constant, the expected within-population coalescence time increases $\propto r^{-1}$ as $r \to 0$.

**Effective parameters** In the completely neutral structured coalescent model for the mainland-island demographic scenario, expected coalescence times depend on $m, N_A$ and $N_B$, where $N_A$ is the population size of the mainland and $N_B$ that of the island. The relevant expressions, written as functions of those three parameters, are

$$\begin{aligned} t_A(m, N_A, N_B) &= N_A \\ t_{AB}(m, N_A, N_B) &= \frac{1}{m} + N_A \\ t_B(m, N_A, N_B) &= \frac{N_B(3 + 2mN_A)}{1 + 2mN_B} \end{aligned} \tag{S5}$$

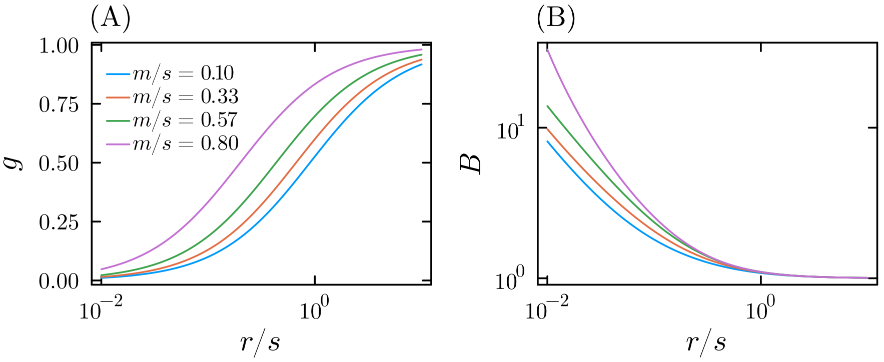

**Figure S1:** Effective parameters $g = \frac{m_e}{m}$ and $B = \frac{N_e}{N}$ for the single barrier model, calculated by solving eq. (S6). We assume the barrier locus is at deterministic migration-selection balance, so that $q = m/s$. $r$ is the recombination rate between the neutral locus at which the effective parameters are evaluated and the barrier locus. In (B) we assume $N_A = 10N_B$.

We now define a set of effective parameters to match the model with a barrier locus to the neutral model. That is, we search for those values of $m, N_A$ and $N_B$ that will yield the expected coalescence times in the presence of a barrier locus when plugged into the neutral expressions eq. (S5). In the model with a barrier locus, we still have $\mathbb{E}[t_A] = N_A$, so we need only two effective parameters, $m_e$ and $N_{B,e} := N_e$ to describe expected coalescence times in the model with a barrier in terms of the neutral expected coalescence times.

$$\begin{aligned}
\mathbb{E}[t_A] &= t_A(m_e, N_A, N_e) = N_A \\
\mathbb{E}[t_{AB}] &= t_{AB}(m_e, N_A, N_e) = \frac{1}{m_e} + N_A \\
\mathbb{E}[t_B] &= t_B(m_e, N_A, N_e) = \frac{N_e(3 + 2m_eN_A)}{1 + 2m_eN_e}
\end{aligned} \tag{S6}$$

Solving for the effective parameters, we find eq. (S3) for $m_e$, as we should, and a complicated expression for $N_e$, the effective size of the island population. These effective parameters are graphed as a function of $r/s$ for different values of $m/s$ in fig. S1.

Notably, when linkage is tight and there is substantial polymorphism at the barrier locus, $N_e$ at the focal neutral locus (as defined by eq. (S6)) can become much larger than the island population size $N_B$, and, consequently, the within-island coalescence time becomes much larger than the neutral prediction (see also fig. S2 A). This is because there is an appreciable probability that the two lineages are on different genetic backgrounds, which, in the case of tight linkage, entails that one of the two lineages is likely to trace back rapidly to the mainland. As a result, the expected within-island coalescence time will be pulled towards $\mathbb{E}[t_{AB}]$, the expected between-population coalescence time.

From the above analysis, it is clear that there is no single effective parameter that suffices to describe genealogies at a neutral locus linked to a barrier locus. For instance, substituting $m_e$ for $m$ in eq. (S5) will not yield the expected within-population coalescence time at a neutral locus linked to a barrier: one needs a second effective parameter (here called $N_e$) to make this work. This is mostly an issue of tight linkage: when $r/s$ is sufficiently large, a single $m_e$ parameter does adequately predict the within-island coalescence time (fig. S2 A). Indeed, when $r \gg s$, sampling lineages on different genetic backgrounds *does not* entail that one of the two traces back rapidly to the mainland, since recombination will likely have reshuffled genetic backgrounds before selection removes a deleterious allele.

Finally, for the case where $N_A \gg N_B$, it is a well known result for the neutral mainland-

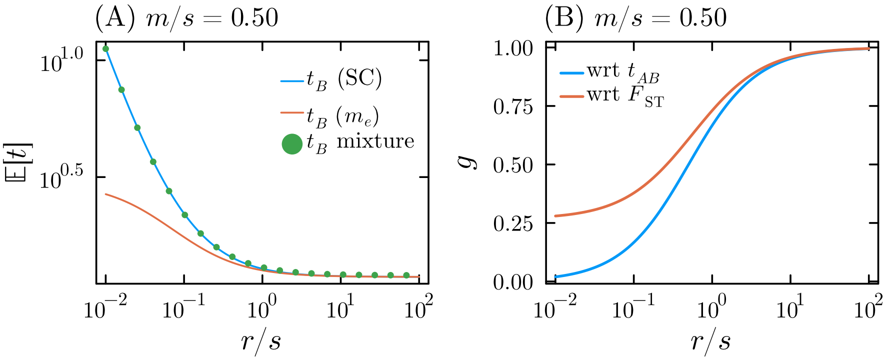

**Figure S2:** (A) Comparison of (1) the structured coalescent prediction for the expected within-island coalescence time ($t_B$, in units of $N_B$) (2) the prediction obtained by plugging $m_e$ into the corresponding expression for the neutral mainland-island model ($t_B^*$) and (3) a mixture model $\mathbb{E}[t_B] = (1 - 2m/s)t_B^* + (2m/s)\mathbb{E}[t_{AB}]$ (see section S1.5). (B) Comparison of the gene flow factor ($g$) defined with respect to the cross-population coalescence time (in blue), and the gff defined with respect to $F_{\mathrm{ST}}$, assuming $N_A \gg N_B$ (i.e. eq. (S7)).

island model that

$$\mathbb{E}[F_{\mathrm{ST}}] = \frac{1}{1 + 2N_B m}$$

Given that $\mathbb{E}[F_{\mathrm{ST}}]$ only depends on the product $N_B m$, one could define an effective parameter with respect to $F_{\mathrm{ST}}$, i.e. define an $m_e$ so that

$$\mathbb{E}[F_{\mathrm{ST}}] = \frac{1}{1 + 2N_B m_e}$$

This yields a gff that can be expressed as

$$g_{F_{\mathrm{ST}}} = \frac{q\left(-M\left(2R+1\right)q + \left(2M+1\right)\left(R+1\right) + q\left(Rp\left(2R+1\right)-1\right)\right)}{2M^2p^2 + 2Mpq\left(R\left(p+1\right)-1\right) + M + Rq\left(2Rpq+1\right)} \tag{S7}$$

where $M = N_B m$ and $R = N_B r$ are the scaled migration and recombination rates. This yields an $m_e$ which is different from eq. (S3), and which does not predict $t_{AB}$ or $t_B$ (fig. S2 B).

#### S1.2 Structured coalescent model for two barrier loci

Let $x_1, x_2, x_3$ and $x_4$ be the frequencies of the `00`, `01`, `10`, and `11` haplotypes for the two selected loci on the island, and let $p_1 = 1 - q_1 = x_1 + x_2$ and $p_2 = 1 - q_2 = x_1 + x_3$. The mainland is assumed to be fixed for the `11` haplotype. Let $r_1$ be the recombination rate between the neutral locus and the first selected locus, and $r_2$ between the neutral locus and the second selected locus. Finally, let $r$ be the recombination rate between the two selected loci.

When the neutral locus is located *in between* the two barrier loci, the backward transition matrix is, to first order in $s$ and $m$,

$$B_{1.2} = \begin{pmatrix} 1 - r_1q_1 - r_2q_2 & \frac{r_2p_2x_2}{x_1} & \frac{r_1p_1x_3}{x_1} & 0 & 0 \\ \frac{r_2q_2x_1}{x_2} & 1 - r_1q_1 - r_2p_2 & 0 & \frac{r_1p_1x_4}{x_2} & 0 \\ \frac{r_1q_1x_1}{x_3} & 0 & 1 - r_1p_1 - r_2q_2 & \frac{r_2p_2x_4}{x_3} & 0 \\ 0 & \frac{q_1r_1x_2}{x_4} & \frac{q_2r_2x_3}{x_4} & 1 - r_1p_1 - r_2p_2 - \frac{m}{x_4} & \frac{m}{x_4} \\ 0 & 0 & 0 & 0 & 1 \end{pmatrix}$$

Where the first four rows and columns correspond to transitions between genetic backgrounds on the island, and the fifth column involves transitions to the mainland background.

When the neutral locus is located on *the left* of the two selected loci, the backward transition matrix is

$$B_{.12} = \begin{pmatrix} 1 - ry_3 - r_1 y_1 & \frac{x_2(p_2 r + r_1 x_1)}{x_1} & r_1 x_3 & r_1 x_4 & 0 \\ \frac{x_1(q_2 r + r_1 x_2)}{x_2} & 1 - ry_4 - r_1 y_2 & r_1 x_3 & r_1 x_4 & 0 \\ r_1 x_1 & r_1 x_2 & 1 - ry_1 - r_1 y_3 & \frac{x_4(p_2 r + r_1 x_3)}{x_3} & 0 \\ r_1 x_1 & r_1 x_2 & \frac{x_3(q_2 r + r_1 x_4)}{x_4} & 1 - ry_2 - r_1 y_4 - \frac{m}{x_4} & \frac{m}{x_4} \\ 0 & 0 & 0 & 0 & 1 \end{pmatrix}$$

where $y_i = 1 - x_i$. The case with the neutral locus on the right follows by symmetry.

**Between-population coalescence times** Given the backward transition matrix $B$, the expected time $t_i$ until absorption into the mainland when in genetic background $i$ on the island is given by a 'first-step analysis' as:

$$\mathbb{E}[t_i] = \frac{1}{\sum_{j \neq i} B_{ij}} \left( 1 + \sum_{j \neq i} B_{ij} \mathbb{E}[t_j] \right) \tag{S8}$$

This is a linear system of four equations in four unknowns that can be readily solved. The expected between-population coalescence time is then $\mathbb{E}[t_{AB}] = \sum_i x_i \mathbb{E}[t_i] + N_A$, where $N_A$ is the size of the mainland population.

**Within-population coalescence times** Similarly, the expected time until coalescence $t_{ij}$ when sampling two lineages in genetic backgrounds $i$ and $j$ (including the mainland background) is

$$\mathbb{E}[t_{ii}] = \frac{1}{\frac{1}{N_i} + \sum_{j \neq i} 2B_{ij}} \left( 1 + \sum_{j \neq i} 2B_{ij} \mathbb{E}[t_{ij}] \right)$$

$$\mathbb{E}[t_{ij}] = \frac{1}{\sum_{k \neq i} B_{ik} + \sum_{k \neq j} B_{jk}} \left( 1 + \sum_{k \neq i} B_{ik} \mathbb{E}[t_{kj}] + \sum_{k \neq j} B_{jk} \mathbb{E}[t_{ik}] \right) \tag{S9}$$

Where $N_i$ is the size of the subpopulation of genetic background $i$. This is a linear system of 15 (i.e. $5 + \frac{5 \times 4}{2}$) equations in 15 unknowns (recall that we include the mainland as a genetic background here, and note that $t_{ij} = t_{ji}$). Let $x_i$ be the frequency of genetic background $i$ on the island, than the expected coalescence time of two lineages sampled from the island is

$$\mathbb{E}[t_B] = \sum_i \sum_j x_i x_j \mathbb{E}[t_{ij}]$$

To calculate coalescence times, we need to calculate the backward matrices, which require knowing the haplotype frequencies. For the two-locus case, equilibrium haplotype frequencies (in a deterministic continuous-time model) were given by Bürger and Akerman (2011). Calculations based on eq. (S8) and eq. (S9) are expected to yield good results whenever $Ns$ is sufficiently large so that haplotype frequencies do not fluctuate much around the corresponding deterministic predictions.

**Effective migration rates** The effective migration rate at the neutral locus is $(\sum_i x_i t_i)^{-1}$, i.e. the inverse of the expected time until absorption in the mainland. If we solve for this symbolically, we obtain very complicated expressions for the gff at a neutral locus in between and on the side of two selected loci that are functions of $q_1$, $q_2$ and $D$, (i.e. the allele frequencies at the selected loci and linkage disequilibrium). If we substitute deterministic predictions (Bürger and Akerman, 2011) for the haplotype frequencies in $B_{1.2}$ and $B_{.12}$ above and take the limit as $m/s \to 0$, we find the following expressions for the gff

$$g_{1.2} = \frac{r_1 r_2}{(r_1 + s_1)(r_2 + s_2)} \tag{S10}$$

$$g_{.12} = \frac{r_1(r_2 + s_1)}{(r_1 + s_1)(r_2 + s_1 + s_2)} \tag{S11}$$

For a neutral locus in between and on the side of two barrier loci respectively. These expressions are identical to those of Aeschbacher and Bürger (2014), there obtained by other means.

#### S1.3 Branching process approximation for the selective $m_e$

For a single selected locus, we find a branching process approximation for the equilibrium allele frequencies by considering a decomposable two-type BP (see e.g. Haccou et al. (2005)) with the following mean matrix

$$M = \begin{pmatrix} e^{-s_1 p_1} & 1 \\ 0 & 1 \end{pmatrix}$$

Such a BP allows us to calculate the total number of descendants of a migrant individual that have inherited the deleterious allele at the selected locus from their migrant ancestor up to any generation $t$ after the migrant arrived on the island. Note that the types in this BP do *not* correspond to the types in the BP that we used for the derivation of the gff at a neutral locus linked to a barrier locus in the main text (i.e. eq. (1)). Specifically, $\mathbb{E}[Z_{t,2}]$, the expected number of type 2 'individuals' in generation $t$, counts the *total* number of descendants (that carry the deleterious allele) that the migrant has had up to generation $t$ (see Haccou et al. (2005) (section 2.3.2) for more on decomposable BPs). Hence, at equilibrium, one should have

$$q_1^* = m \lim_{t\to\infty} \mathbb{E}[Z_{t,2}] \tag{S12}$$

Denoting by $\nu_1$ the leading eigenvector of $M$. This equilibrium allele frequency can be found to be

$$q_1^* = [m \quad 0]\nu_1 \approx \frac{m}{s_1 p_1} \tag{S13}$$

this overpredicts the deleterious allele frequency whenever there is partial divergence, due to violation of the branching property.

For two selected loci, a similar analysis yields

$$M = \begin{pmatrix} e^{-s_1p_1 - s_2p_2}(1-r) & e^{-s_1p_1-s_2p_2} r & 1 \\ 0 & e^{-s_1p_1} & 1 \\ 0 & 0 & 1 \end{pmatrix}$$

$$\tilde{q}_1 = [m \quad 0 \quad 0]\nu_1 \approx m \frac{r_1 + s_1 p_1}{s_1 p_1 (r_1 + s_1 p_1 + s_2 p_2)}$$

Combining this with eq. (S13), we find

$$g := \frac{\tilde{q}_1}{q_1^*} \approx \frac{r_1 + s_1 p_1}{r_1 + s_2 p_2 + s_1 p_1}$$

For the three-locus case we consider both the case with focal locus in the middle and the focal locus on the edge. In the latter case we find

$$g_{.12} \approx \frac{(r_1 + s_1 p_1)(r_2 + s_1 p_1 + s_2 p_2 s_2)}{(r_1 + s_1 p_1 + s_2 p_2)(r_2 + s_1 p_1 + s_2 p_2 + s_3 p_3)} \tag{S14}$$

whereas for the locus in the middle, we obtain

$$\begin{aligned} g_{1.2} \approx & \frac{(r_1 + s_1 p_1)(r_2 + s_2 p_2)}{(r_1 + s_1 p_1 + s_2 p_2)(r_2 + s_2 p_2 + s_3 p_3)} \\ & + \frac{s_1 p_1 s_2 p_2 s_3 p_3}{(r_1 + s_1 p_1 + s_2 p_2)(r_2 + s_2 p_2 + s_3 p_3)(r_1 + r_2 + s_1 p_1 + s_2 p_2 + s_3 p_3)} \end{aligned}$$

In the multi-locus generalization we ignore the second term.

#### S1.4 Gene flow factor and fixed-point iteration for bidirectional migration

We can use the mainland-island model predictions to obtain an approximation for the more complicated case with bidirectional migration. Our approach decomposes the bidirectional model into two unidirectional mainland-island models, assuming there is no ‘back-migration’, i.e. migrants are always ’resident’ individuals.

Denote the equilibrium allele frequencies in population $A$ and $B$ at divergently selected locus $i \in [1..L]$ by $p_{A,i}$ and $p_{B,i}$. The allele (‘0’) at frequency $p_{A,i}$ in population $A$ is assumed to be beneficial in population $B$ and deleterious in population $A$, whereas the allele (‘1’) at frequency $q_{A,i}$ in $A$ is assumed to be beneficial in $A$ and deleterious in $B$. At equilibrium, we assume that a migrant from $B$ entering $A$ has a genotype that is randomly drawn from the $B$ population at Hardy-Weinberg and linkage equilibrium (HWLE), and similarly for a migrant from $A$ entering population $B$. The gffs for $B \to A$ and $A \to B$ migration can then be written, respectively, as

$$\begin{aligned} g_k^{BA} &\approx \exp\left( -\sum_{i \in L_l} \frac{a_i \Delta_i}{r_i + a_k \Delta_k + \sum_{j<i} a_j \Delta_j} - \sum_{i \in L_r} \frac{a_i \Delta_i}{r_i + a_k \Delta_k + \sum_{j<i} a_j \Delta_j} \right) \\ g_k^{AB} &\approx \exp\left( -\sum_{i \in L_l} \frac{b_i \Delta_i}{r_i + b_k \Delta_k + \sum_{j<i} b_j \Delta_j} - \sum_{i \in L_r} \frac{b_i \Delta_i}{r_i + b_k \Delta_k + \sum_{j<i} b_j \Delta_j} \right) \end{aligned}$$

where $\Delta_i = p_{B,i} - p_{A,i}$ is the allele frequency divergence at locus $i$, $a_i$ is the selection coefficient against the allele with allele frequency $p_{A,i}$ in population $A$, and $b_i$ is the selection coefficient against the allele with frequency $q_{B,i}$ in population $B$.

Starting from complete divergence ($p_{A,i}^{(0)} = 0$ and $p_{B,i}^{(0)} = 1$ for $i = 1, \ldots, L$), we can use a similar fixed-point iteration as before. Specifically, given the allele frequencies $p_{A,i}^{(k)}, p_{B,i}^{(k)}$ obtained in iteration $k$, we execute the following steps

1. calculate $g_j^{BA}$ and $g_j^{AB}$ for $j = 1, \ldots, L$
2. calculate $p_{A,j}^{(k+1)} = \int_0^1 p\phi\left(p|N, u, s_j, m_{BA}, g_j^{BA}, p_{B,j}^{(k)}\right) dp$ where

$$\phi(p|N, u, s, m, g, p_B) = \frac{p^{2N(u+mp_B)-1)} q^{2N(u+m(1-p_B))-1} e^{-Nsq}}{\int_0^1 p^{2N(u+mp_B)-1} q^{2N(u+m(1-p_B))-1} e^{-Nsq} dq} \tag{S15}$$

3. calculate $p_{B,j}^{(k+1)} = \int_0^1 p\phi\left(p|N, u, s_j, m_{AB}, g_j^{AB}, p_{A,j}^{(k)}\right) dp$

and we iterate until convergence.

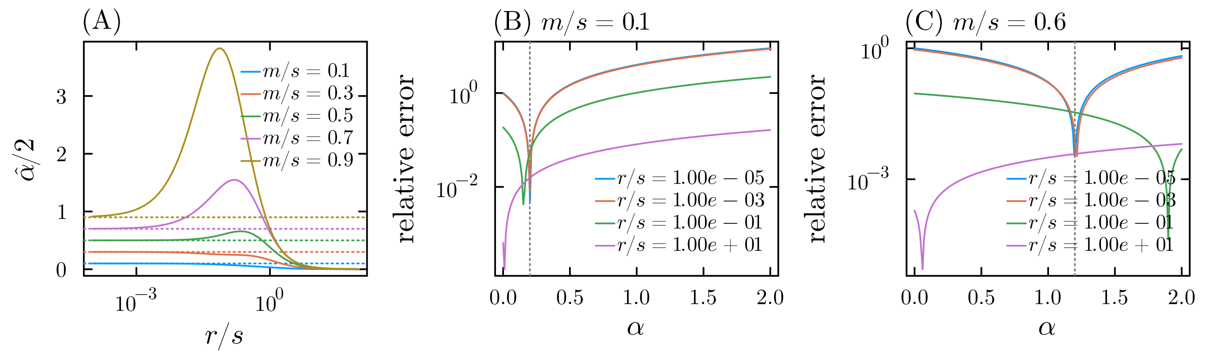

**Figure S3:** (A) Optimal $\alpha$ ($\hat{\alpha}$ graphed as $\hat{\alpha}/2$) as a function of $r/s$ for various values of $m/s$. The horizontal lines mark the value of $m/s$. (B) Relative error of the mixture approximation for the within-island coalescence time in the single barrier model, i.e. $|\mathbb{E}_{\text{mix}}[t_B] - \mathbb{E}_{\text{SC}}[t_B]|/\mathbb{E}_{\text{SC}}[t_B]$ as a function of the mixture proportion $h$ for various degrees of linkage. The vertical line marks $\alpha = m/s$. We assume $m/s = 0.1, s = 0.02, N_A = N_B = 1000$. (C) as in (B) but for $m/s = 0.6$.

#### S1.5 Within-population coalescence time approximation

In section S1.1 we showed that $m_e$ does not suffice to predict the within-island coalescence time, and that a second effective parameter is needed in order to predict coalescence times in the presence of a single barrier locus by means of the neutral structured coalescent. While we could calculate this second effective parameter for the single-barrier case, it is not clear how to do this in the general polygenic case.

However, we found that, to a first approximation, a two-component mixture model can be used to approximate the expected within-population coalescence time, i.e.

$$\mathbb{E}[t_B] \approx (1-\alpha)t_B^* + \alpha t_{AB}^* \tag{S16}$$

where $t_B^*$ and $t_{AB}^*$ are the neutral mainland-island predictions with $m_e$ substituted for $m$, i.e.

$$t_{AB}^* = \frac{1}{m_e} + N_A \tag{S17}$$

$$t_B^* = \frac{N_B(3 + 2m_e N_A)}{1 + 2m_e N_B} \tag{S18}$$

Heuristically, $\alpha$ can be thought of as the probability that one of the two sampled lineages has recent migrant ancestry and moves into the mainland on a very short timescale compared to $1/m_e$. Hence, conditional on sampling one lineage with recent migrant ancestry and one resident lineage, the coalescence time should be close to $t_{AB}^*$, whereas the coalescence time should be close to $t_B^*$ otherwise (this ignores the possibility of sampling two lineages with recent migrant ancestry).

**Mixture approximation for the single barrier case** In the single-barrier case, we find that $\alpha = 2q = 2m/s$ minimizes the error between the full structured coalescent prediction (i.e. using eq. (S4) and eq. (S9)) and the mixture apporximation (eq. (S16)) as $r/s \to 0$, whereas the optimal $\alpha$ vanishes as $r/s \to \infty$ (figs. S3 and S4). For small $m/s$, this supports our heuristic interpretation above: the probability that one of the two sampled lineages is a recent descendant from a migrant is roughly $2m/s$, in which case the coalescence time should be close to $t_{AB}^*$. Surprisingly, $\alpha = 2m/s$ minimizes the error even when $m/s$ is not small (fig. S3). Even more surpisingly, this extends beyond $m/s = 0.5$ (fig. S3C), so that $\alpha > 1$ and hence can no longer be interpreted as a mixture proportion.

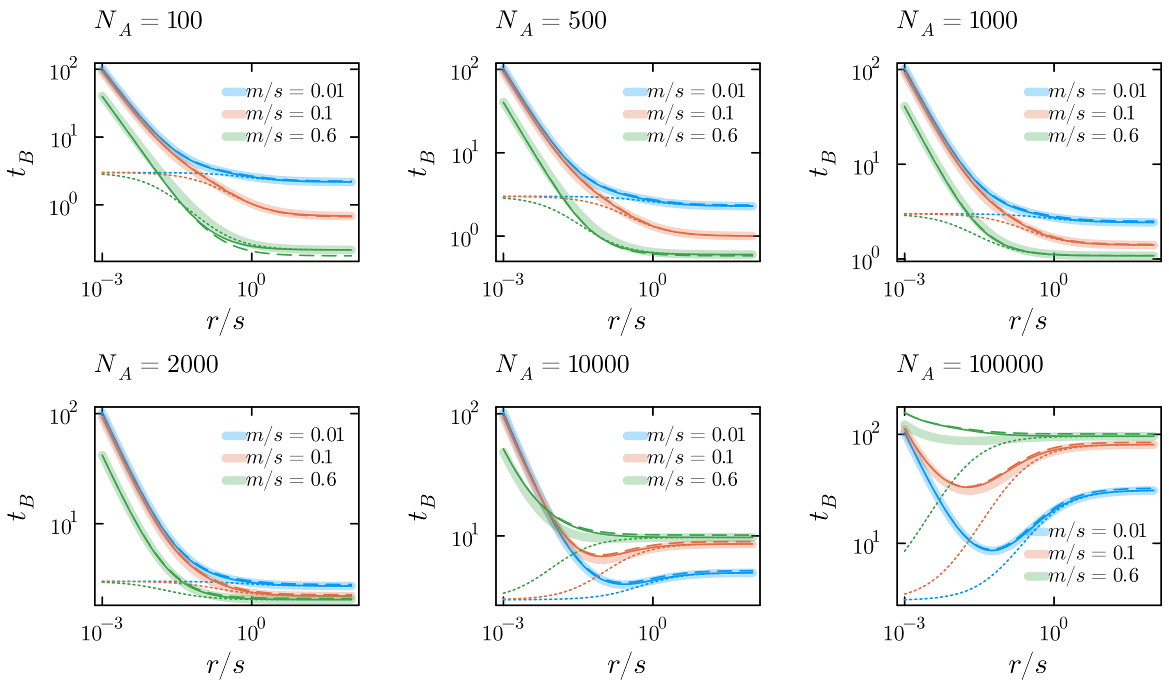

**Figure S4:** Comparison of the structured coalescent prediction for the expected within-island coalescence time ($t_B$) (thick transparent solid lines) against the naive $m_e$ prediction ($t_B^*$, eq. (S18), dotted lines), the mixture model approximation with $\alpha = 2q$ (eq. (S16), dashed lines) and the mixture model approximation with $\alpha = 2(1-g)q$ (eq. (S16) and eq. (S19), solid lines) for the single barrier case for different migration rates ($m/s$) and mainland population sizes ($N_A$). We assume $s = 0.02$ and an island population size $N_B = 1000$. Time is measured in $N_B$ generations.

From the above analysis, we see that the mixture proportion should depend on genomic location ($r/s$): for a neutral locus closely linked to a barrier, sampling an individual with recent migrant ancestry entails that the neutral lineage is likely to trace back to the mainland rapidly, whereas for a neutral locus that is not linked to the barrier this is not quite the case. From the BP model (see eq. (1) in the main text), one can obtain an approximation to the proportion of 'nonresidents' ($\pi_{nr}$, the proportion of individuals that have inherited migrant alleles at both the neutral and barrier locus) at equilibrium, i.e.

$$\pi_{nr} = \frac{m}{r + sp} = \frac{m_e}{r} = \frac{q}{\frac{r}{s} + p} = (1-g)\frac{q}{p}$$

we find that using

$$\alpha = 2\pi_{nr}p = 2\frac{mp}{r + sp} = 2(1-g)q \qquad \text{(S19)}$$

we obtain a very good approximation to $\mathbb{E}[t_B]$ using the mixture approximation (fig. S4). Equation (S19) yields the same results as the structured coalescent prediction in both the limit of loose and tight linkage, i.e. $\alpha \to 0$ as $s/r \to 0$, and $\alpha \to 2q$ as $r/s \to 0$. As expected, we find that the error in the mixture approximation increases with $m/s$, and that this is more pronounced when the size of the mainland and island differ appreciably (fig. S4).

**Mixture approximation in the polygenic case** Empirically, we found that eq. (S16) can predict within-population coalescence times remarkably well also in the polygenic regime. This is illustrated in figs. S19 to S22, where we fitted eq. (S16) by finding the $\alpha$ that minimizes the sum of squared deviations between observed within-population coalescence times in individual-based simulations and predictions. Note that this fits a single $\alpha$ for the entire genome, irrespective of genomic location.

In order to *a priori* predict the mixture proportion $\alpha$, we continue on the basis of our heuristic interpretation, and think of $\alpha = 2h$ as the probability to sample a lineage that has recent migrant ancestry, or equivalently, consider $h$ as the proportion of recent migrant ancestry at any genomic position in the population as a whole. As in our empirical fitting procedure, we assume that that the mixture proportion does not depend on genomic location, and that heterogeneity in coalescence times is fully captured by the heterogeneity in $t^*_{AB}$ and $t^*_B$ (which derives from heterogeneity in $m_e$). In the polygenic regime this can be motivated as follows: in the first couple of generations after migration, purging of migrant ancestry proceeds by eliminating large blocks of genome, corresponding to a loss of about 50% of migrant ancestry per generation. When deleterious alleles are spread more or less uniformly across the genome, the probability to survive this initial phase of rapid purging should be roughly independent of genomic location. Only in a later stage does the detailed linkage structure become important in determining the long-term survival of particular blocks of genome.

**Predicting the proportion of recent migrant ancestry $h$** Our goal is to approximately predict the probability that a sampled lineage at some neutral locus in the genome descends from a migrant in the recent past. At equilibrium, the proportion of $k$th generation descendants of migrants is approximately

$$f_k = 2^k m W_0 W_1 \dots W_{k-1} + O(m^2) \tag{S20}$$

where $k = 1$ refers to F1s, $k = 2$ to BC1s, *etc.* and $W_k$ is the mean relative fitness of $k$th generation migrant descendants, *relative to residents.* Let $h_k$ be the proportion of migrant ancestry in the population as a whole that is in $k$th generation descendants. The proportion of migrant ancestry in the population as a whole is then $h = \sum_{k=1}^{c} h_k$, where $c$ is some upper bound on what is counted as a migrant (i.e. non-resident) generation.

When linkage between selected loci is loose on average, an F1 carries about 50% migrant ancestry and 50% of the deleterious alleles carried by its migrant parent, a BC1 25% *etc.*, so that one obtains

$$h_k = \frac{f_k}{2^k} = m W_0 W_1 \dots W_{k-1} \tag{S21}$$

Assuming that we know allele frequencies in the island population and that the mainland is fixed for the locally deleterious alleles, we have the following recursion

$$h_{k+1} = h_k W_k \approx \exp\left(-\frac{\sum_i^L s_i p_i}{2^k}\right) h_k \tag{S22}$$

with $h_0 = m$. This approximation ignores the variance in fitness generated by segregation. Note that since $W_k \to 1$ with increasing $k$, $h = \sum_{k=1}^{c} h_k$ will not converge as $c \to \infty$, hence a calculation of $h$ will depend on $c$, i.e. on our truncation of what counts as a 'recent' descendant from a migrant (non-resident).

When linkage between selected loci is appreciable, purging of migrant ancestry will be more efficient (i.e. $h_k$ will be smaller than its unlinked approximation), as the variance in fitness among descendants is increased. To account for this, we use an approximation due to Veller et al. (2023), adapted to our multilocus model:

$$h_{k+1} = \exp\left(-\overline{(1-r)^k} \sum_i^L s_i p_i\right) h_k \tag{S23}$$

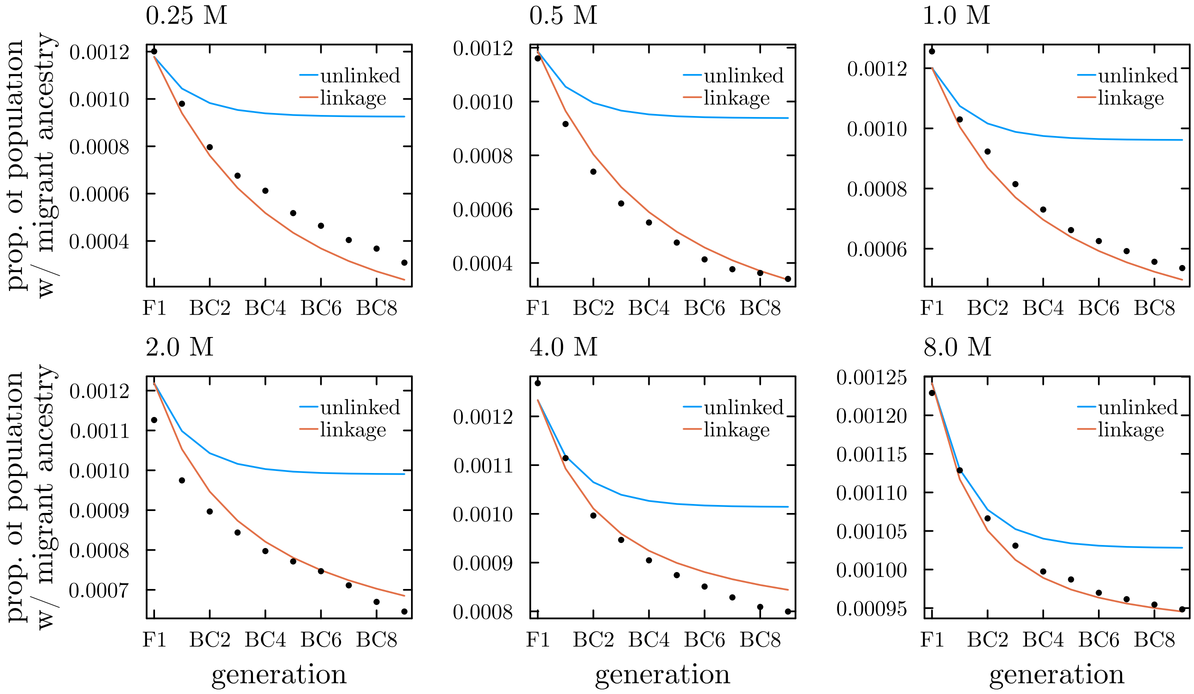

**Figure S5:** Predictions for the proportion of the island population with recent migrant ancestry. Here we assume $L\bar{s} = 0.25, L = 50$ and $N_B\bar{s} = 5$, with the $s_i$ exponentially distributed with mean $\bar{s}$. Results are shown for six different total map lengths. The dots show results from individual-based simulations, where we first simulate the population until equilibrium is approximately reached, and then simulate for 5000 additional generations, tracing back the proportion of migrant ancestry in each generation that is found in F1, BC1, ..., BC9 individuals. The blue line shows predictions based on the unlinked approximation (eq. (S22)), whereas the red line shows predictions based on eq. (S23).

Here the term $\overline{(1-r)^k}$ is an average over the $L \times (L-1)/2$ pairwise recombination rates, i.e. $2\sum_i \sum_{j<i}(1-r_{ij})^k/L(L-1)$. Note that the unlinked approximation appears as a special case. This approximation appears to yield reasonably good predictions when compared against simulations (fig. S5). In our mixture approximation, we use eq. (S23) to calculate $h$, choosing $c = 20$ as an arbitrary threshold (This threshold appears to yield fairly good predictions in figs. S19 to S22). Importantly, eq. (S23) relies on our ability to predict allele frequencies ($p_i$) at divergently selected loci.

### S2 Supplementary figures

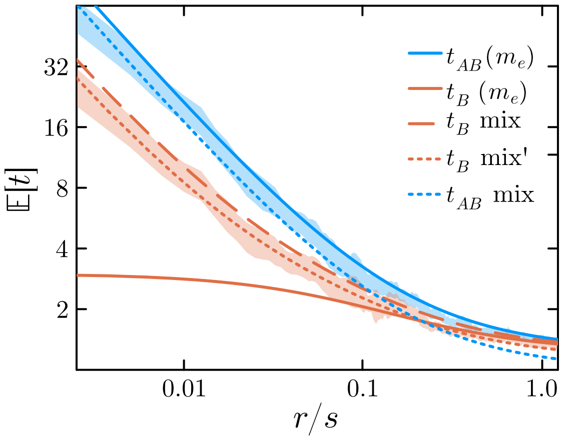

**Figure S6:** Various $m_e$-based approximations to the expected within-island and between-population pairwise coalescence times as a function of degree of linkage to the barrier locus ($r/s$). The colored bands show the same results as in fig. 2. $t^*_{AB}$ and $t_B(m_e)$ are the predictions based on plugging in $m_e$ for $m$ in the neutral prediction. $t_B$ mix uses eq. (S16) with $\alpha = 2q = 2m/s$. $t_{AB}$ mix assumes $\mathbb{E}[t_{AB}] = (1-q)t^*_{AB}$ while $t_B$ mix' assumes $\mathbb{E}[t_B] = (1-2q)t_B(m_e) + 2q(1-q)t^*_{AB}$.

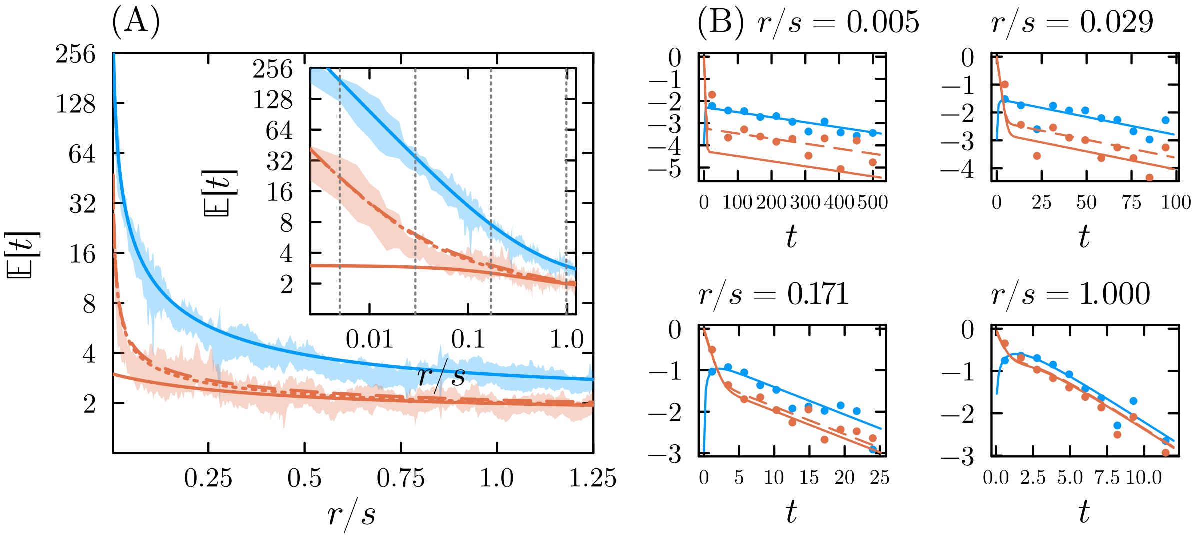

**Figure S7:** As in fig. 2 but with $m/s = 0.05$.

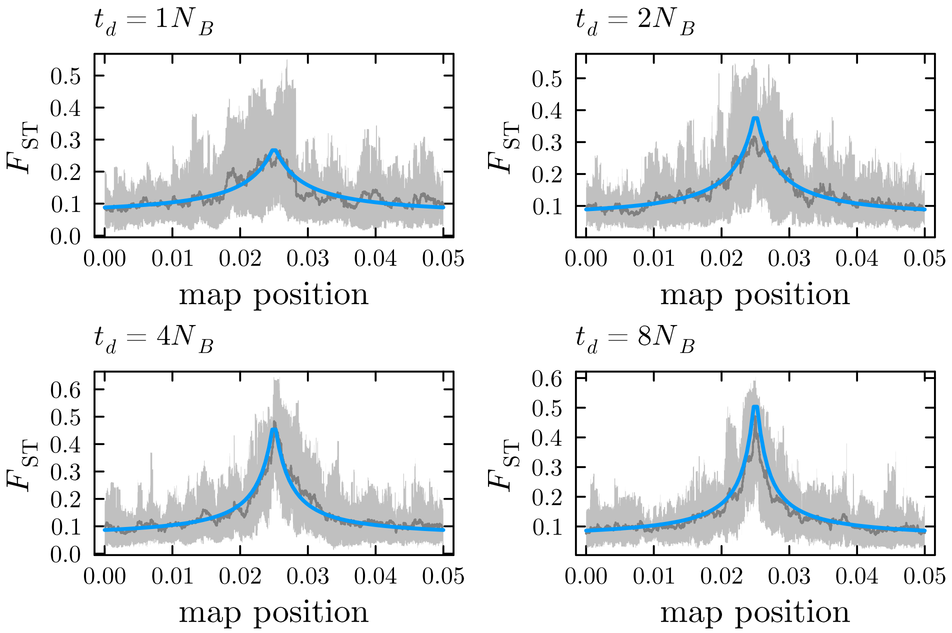

**Figure S8:** $F_{ST}$ patterns near a single barrier locus ($N_B = N_A = 1000, N_B s = 10, m/s = 0.2$) when the island population branched off from the mainland population $t_d$ generations in the past, after which divergent selection started immediately and the population was at migration-selection balance instantaneously. The gray shaded area shows 90% density intervals for the mean $F_{ST}$ across the full population sample, with the gray line showing the mean. The blue line shows the $m_e$ based prediction (using the `PhaseGen` library).

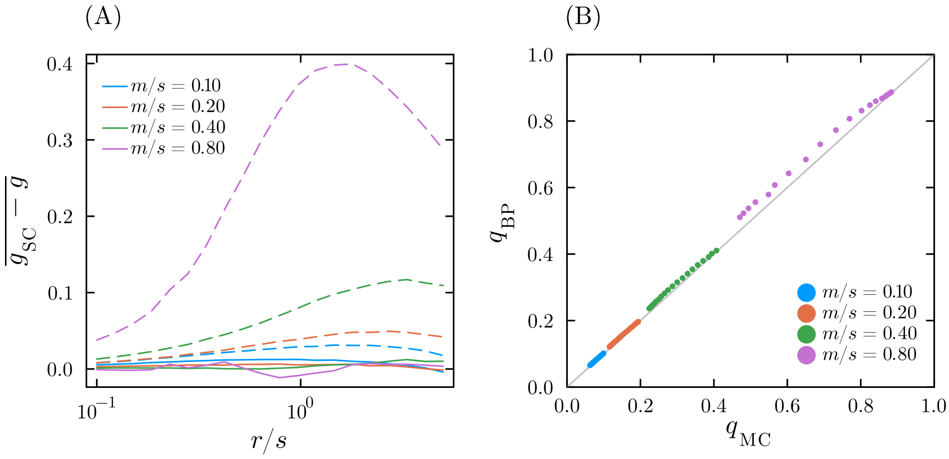

**Figure S9:** (A) Comparison of the Aeschbacher and Bürger (2014) (dashed line) and BP (solid line) predictions of the gene flow factor $g$ against the structured coalescent prediction $g_{SC}$ for the two-barrier model, where $r$ is the recombination rate between the barriers, and $s$ the selection coefficient against migrant alleles. We show, for different values of $m/s$, and for each value of $r/s$, the average difference between $g$ and $g_{SC}$ across the genetic map, where the total map length is assumed to be $5r$. (B) Barrier allele frequencies ($q_{BP}$) predicted by the BP approximation, compared against Monte Carlo estimates ($q_{MC}$) from forward-in-time simulations. The latter predictions were used for the structured coalescent calculations in (A) (recall that these depend on $q$ and $D$).

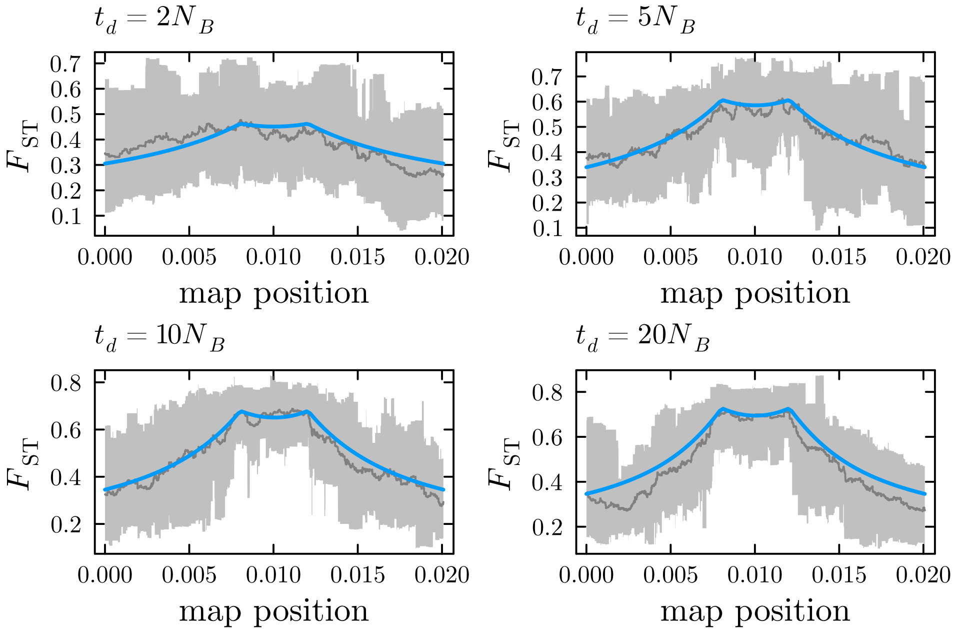

**Figure S10:** Expected between-population coalescence time and 90% interval for a model with two barrier loci where the time since divergent selection is $t_d$ generations before the present. As in fig. S8. We assume $N_A = N_B = 500, N_B s = 10, m/s = 0.2, r/s = 0.2$.

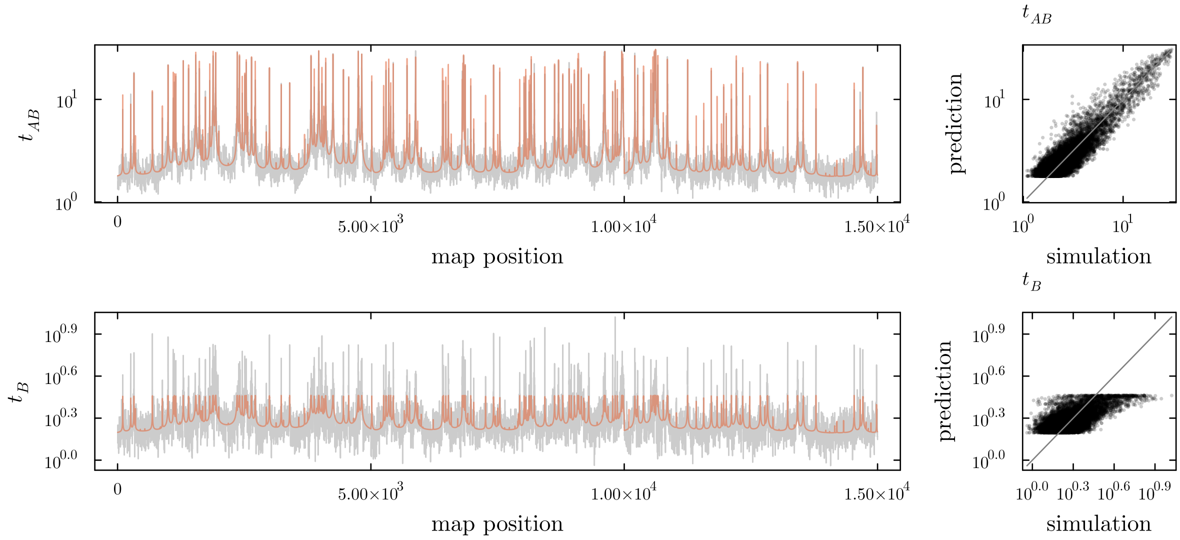

**Figure S11:** Between population ($t_{AB}$) and within-island ($t_B$) pairwise coalescence times (in units of $N_B$ generations) associated with fig. 5. The red lines in the left plot show predicted values using the $m_e$ theory.

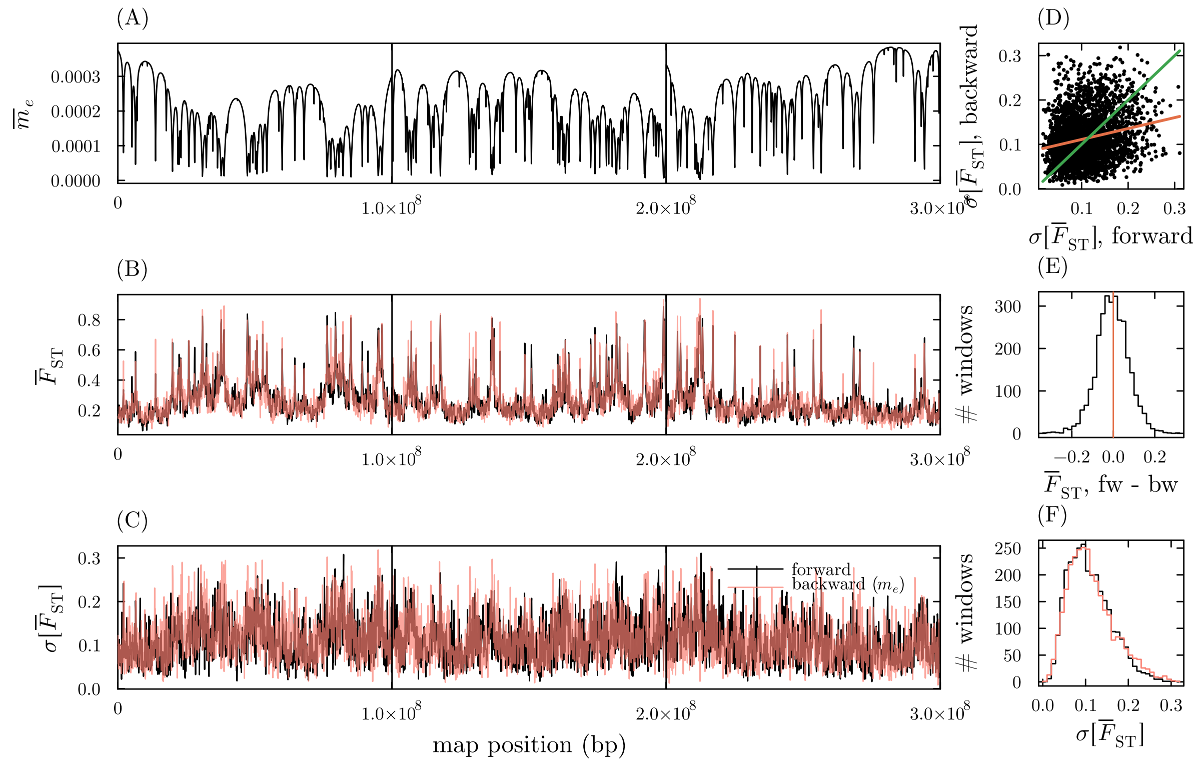

**Figure S12:** (A) Mean $m_e$ in 100kb windows for the simulation shown in fig. 5. (B) Mean $F_{\text{ST}}$ in 100kb windows estimated from forward simulations (black) and backward simulations we simulate SNP data in a window-by-window fashion usingthe neutral structured coalescent with the mean $m_e$ for the relevant window substitued for $m$. We assume $\mu = 10^{-8}$ (equal to the per-base recombination rate). (C) As in (B) but showing the standard deviation of the mean $F_{\text{ST}}$ in windows. (D) Scatter plot for standard deviation of mean $F_{\text{ST}}$ across windows as estimated from forward and backward simulations (each dot is a window). The red line shows the associated linear regression. (E) Difference in mean $F_{\text{ST}}$ in windows estimated from forward and backward simulations. (F) Histograms for the standard deviation of window $F_{\text{ST}}$ for both the forward and backward simulations. All results are based on 5 replicate simulations, taking a sample of 50 haploid genomes in each population.

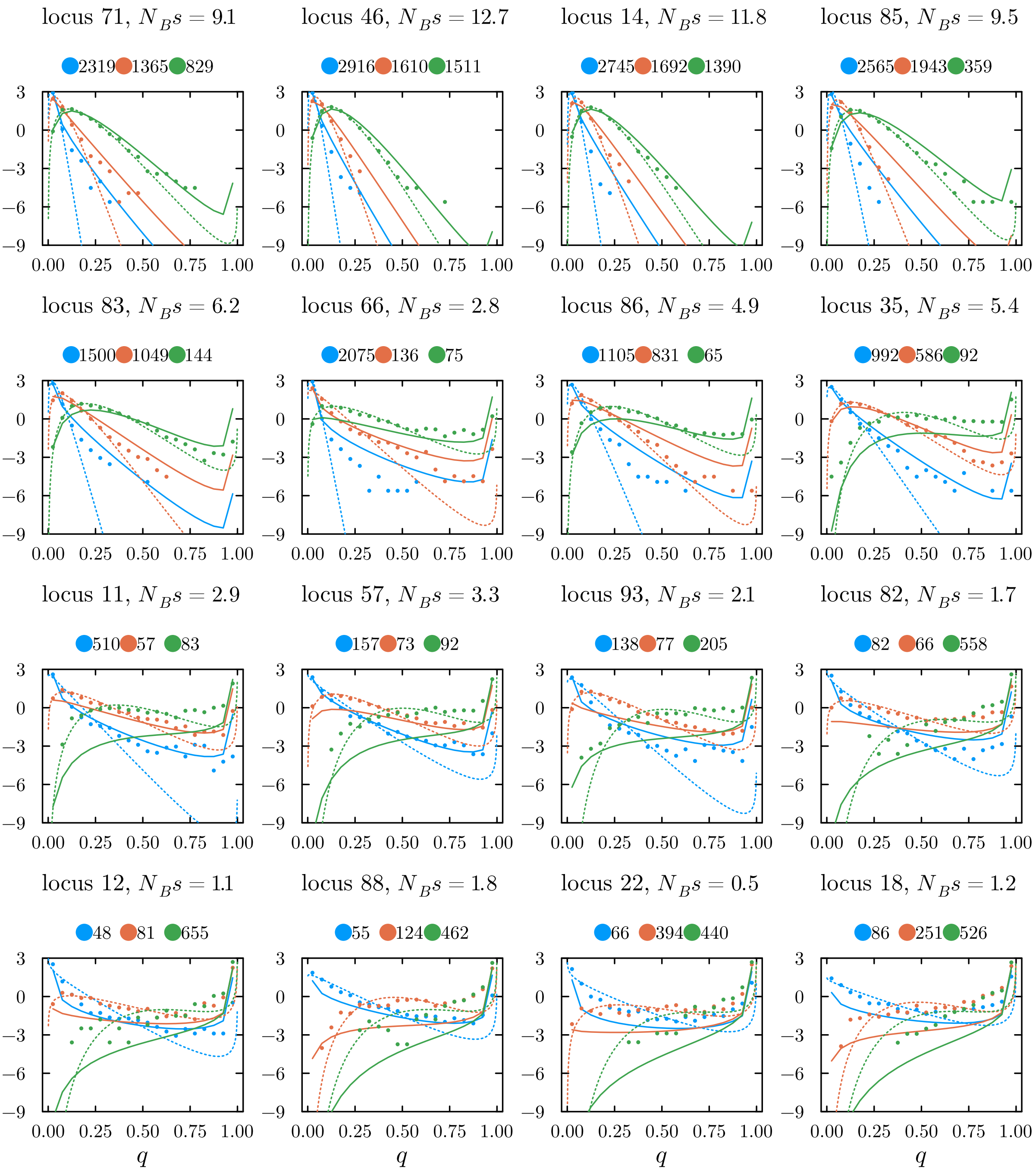

**Figure S13:** Allele frequency distributions for 16 (out of 100) loci in simulation fig. 7, ordered from low to high predicted deleterious allele frequency ($q$) on the island (i.e. high to low predicted adaptive divergence). The solid lines show predicted allele frequency distributions using Wright's density with $m_e$ substituted for $m$, whereas the dots show estimates from forward simulations (the same set of replicates shown in fig. 7. The dotted lines show the best fitting Wright's density with both a free $s_e$ and $m_e$ parameter (using maximum likelihood). The $y$-axis shows the log density. Blue, orange and green show results for map lengths of 25, 50 and 100cM respectively. The numbers above each plot show the effective sample size (estimated effective number of independent samples) for the allele frequencies estimated from the individual-based simulations.

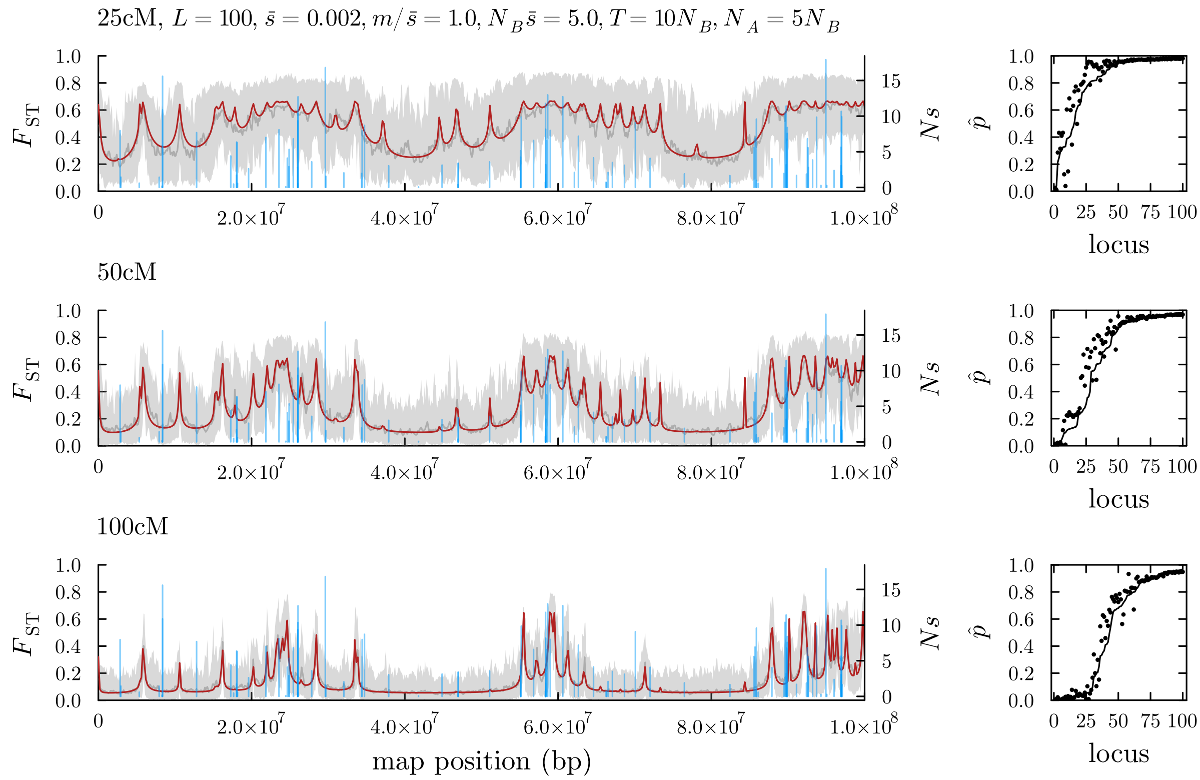

**Figure S14:** As in fig. 7 but with a 'clustered' architecture. We assume that the vector of distances between consecutive loci are distributed as $\sim C \times X$ where $C$ is the map length and $X \sim \text{Dirichlet}(\alpha, L+1)$, where $\alpha$ is the concentration parameter ($\alpha = 1$ amounts to the uniform case, $\alpha > 1$ yields loci that are more evenly spaced on average than the uniform case, and $\alpha < 1$ yields loci that tend to more clustered than the uniform case). Here we assume $\alpha = 0.2$. Other parameters are as in fig. 7 and are listed in the figure titles.

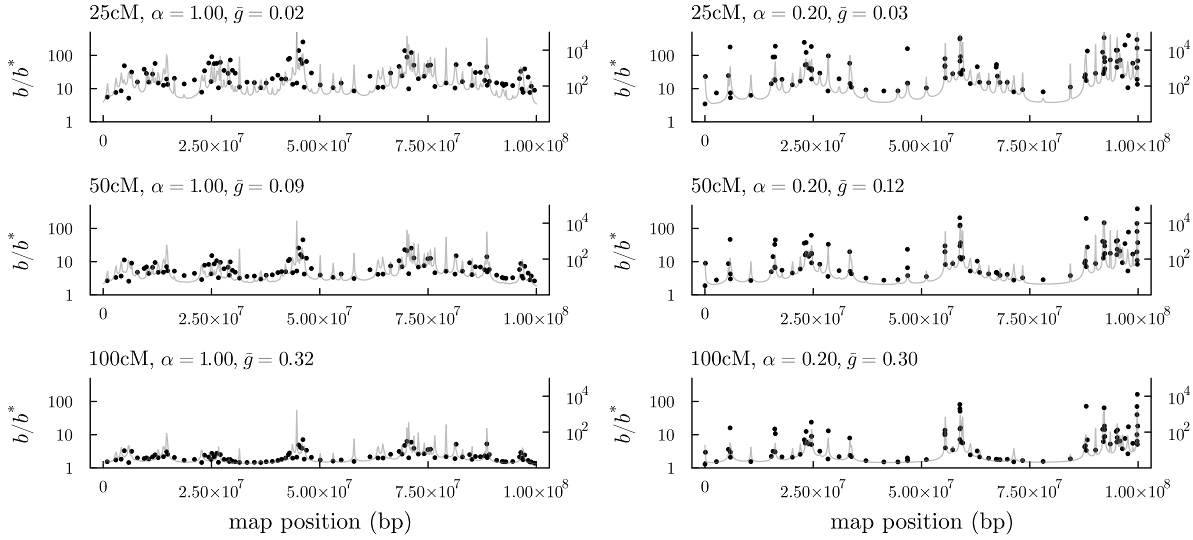

**Figure S15:** Divergence hitchhiking measure $b/b^*$ (see main text and fig. 6) for selected loci in the example simulations of fig. 7 (left column) and fig. S14 (right column). The gray line shows the neutral barrier strength across the chromosome. Values of $\bar{g}$ in the plot titles refer to the genome-wide average neutral gff.

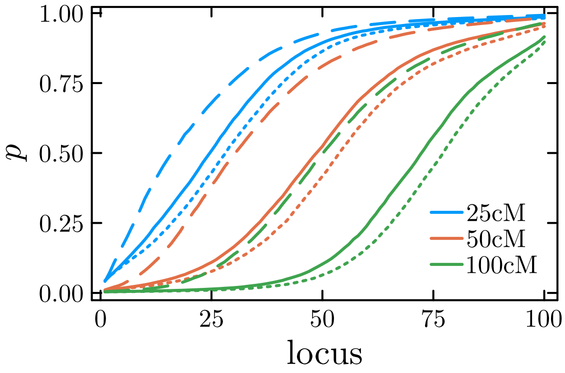

**Figure S16:** Average ranked predicted equilibrium beneficial allele frequencies for genetic architectures consisting of $L = 100$ loci. We assume that the vector of distances between consecutive loci are distributed as $\sim C \times X$ where $C$ is the map length and $X \sim \text{Dirichlet}(\alpha, L+1)$, see fig. S14. The dotted, solid and dashed lines show averages based on 200 random architectures simulated assuming $\alpha = 5, 1, 0.2$ respectively (i.e. for each of the 200 replicates, we draw a random genetic architecture, predict the beneficial allele frequencies using eq. (8) and the fixed point iteration, and rank the predicted allele frequencies). Other parameters are as in fig. S14.

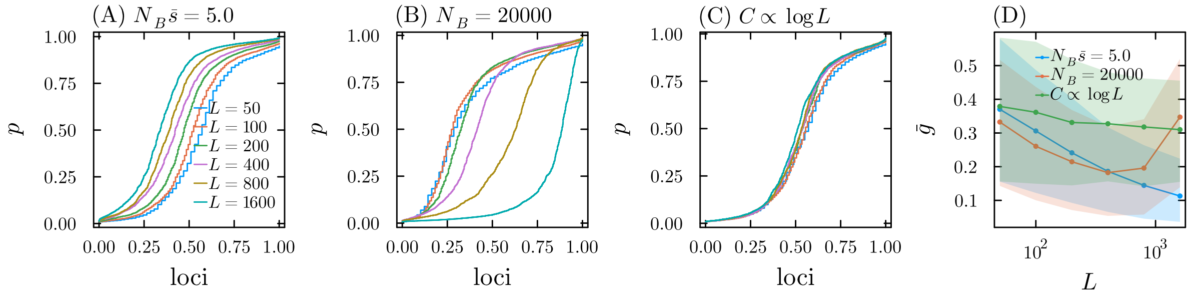

**Figure S17:** The effect of increasing polygenicity. We consider the effects of increasing the number of divergently selected loci $L$ on a single chromosome of map length $C$, keeping $L\bar{s}$ fixed at 0.1, where the loci have exponentially distributed fitness effects with mean $\bar{s}$. (A) Proportion of loci having the predicted locally beneficial allele frequency $\leq p$ when we keep $N_B\bar{s} = 5$ constant as $L$ increases, with $C = 50$cM. (B) As in (A) but where we keep $N_B = 20000$ constant as $L$ increases. (C) As in (A) but where we scale the map length by $\log L$, i.e. when there are $L$ loci, the map length is set to $50\frac{\log L}{\log L_0}$cM, where $L_0 = 50$ (the minimum number of loci considered). This scaling results in an approximately constant harmonic mean recombination rate among the $L$ selected loci as $L$ increases. (D) The average and 90% interval of predicted gff values at selected loci for the barrier architectures examined in (A) (blue) and (B) (orange) and (C) (green). All results are based on sampling $n$ replicate random architectures with $L$ loci uniformly spread across the chromosome, with $n$ chosen such that $nL = 1600$.

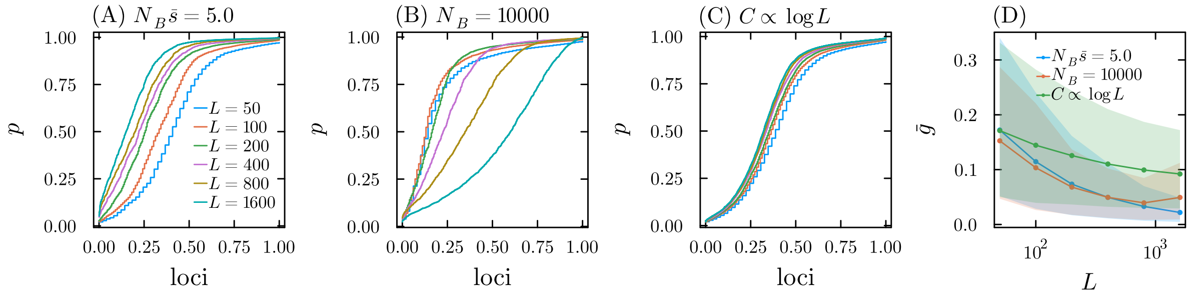

**Figure S18:** As in fig. S17, but assuming $L\bar{s} = 0.2$.

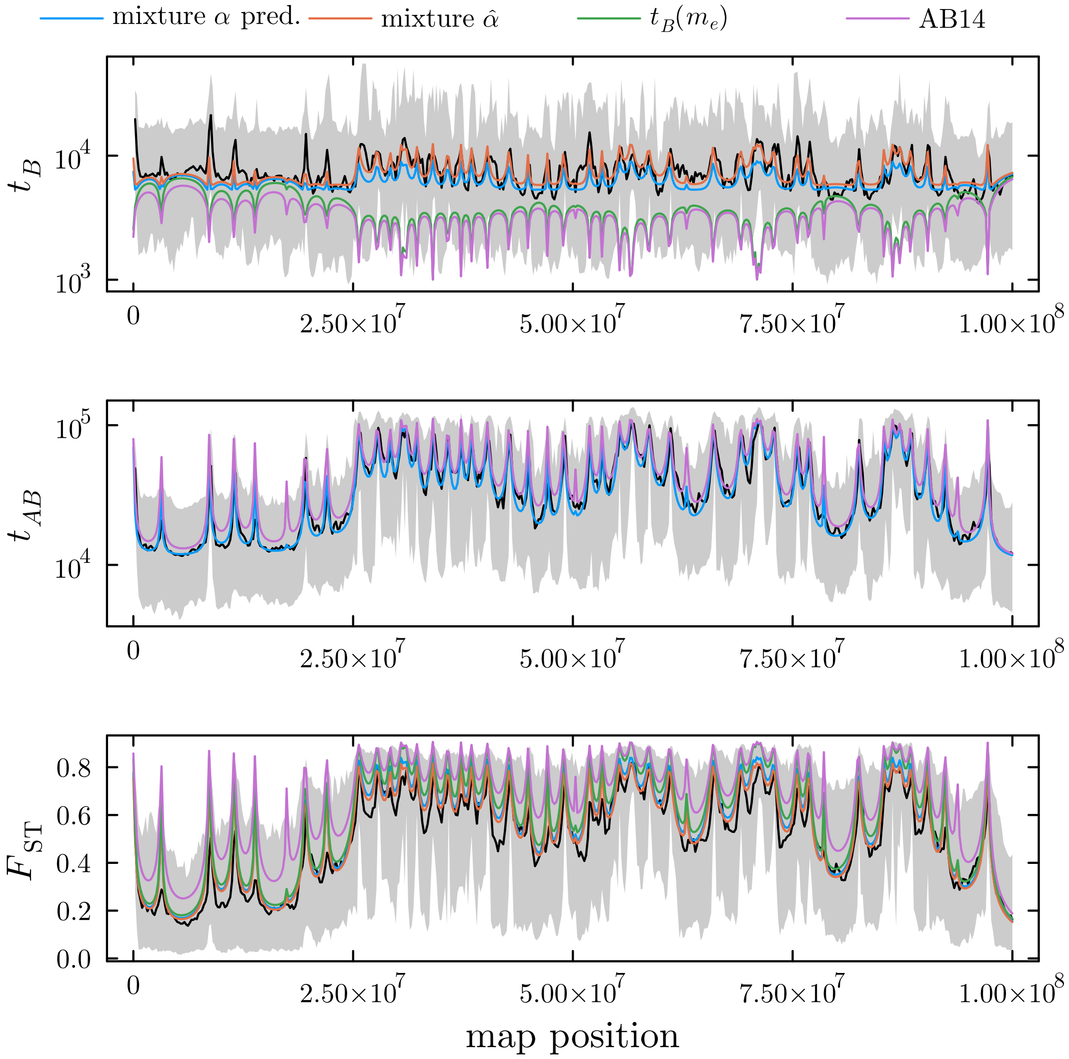

**Figure S19:** Predictions for $\mathbb{E}[t_B]$, $\mathbb{E}[t_{AB}]$ and $F_{\text{ST}}$ compared against results from individual-based simulations. Here we assume a 25cM/100Mb map, and $L\bar{s} = 0.25, L = 50, N_B\bar{s} = 5, m/\bar{s} = 1.5$. The gray line shows the mean estimated from individual-based simulations, with the gray band the associated 95% interval. The blue line shows the mixture approximation using the *a priori* predicted value for $\alpha$ (calculated based on the predicted proportion of migrant ancestry in the first 20 backcross generations). The orange line shows the best fitting mixture approximation ($\hat{\alpha} = 0.1$). The green line shows the naive approximation based on plugging $m_e$ into the neutral structured coalescent prediction for the within-population coalescence time). The purple line shows the prediction based on the Aeschbacher and Bürger (2014) theory. Note that the first three are all equivalent when it comes to predicting the cross-population coalescence time $t_{AB}$ (i.e. $\mathbb{E}[t_{AB}] = 1/m_e + N_A$).

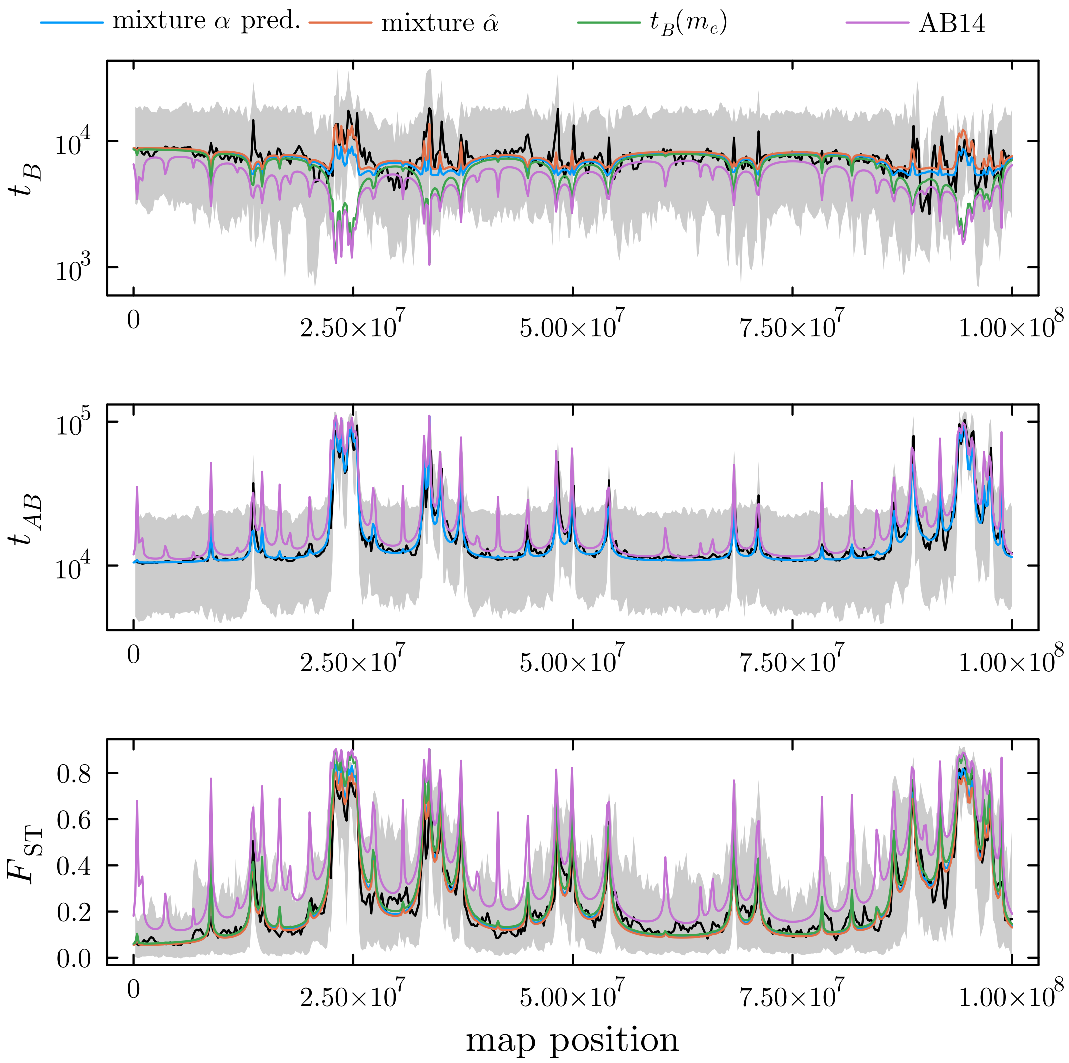

**Figure S20:** As in fig. S19 but with parameters: 25cM, $L\bar{s} = 0.25, L = 50, N_B\bar{s} = 5, m/\bar{s} = 1, \hat{\alpha} = 0.12$.

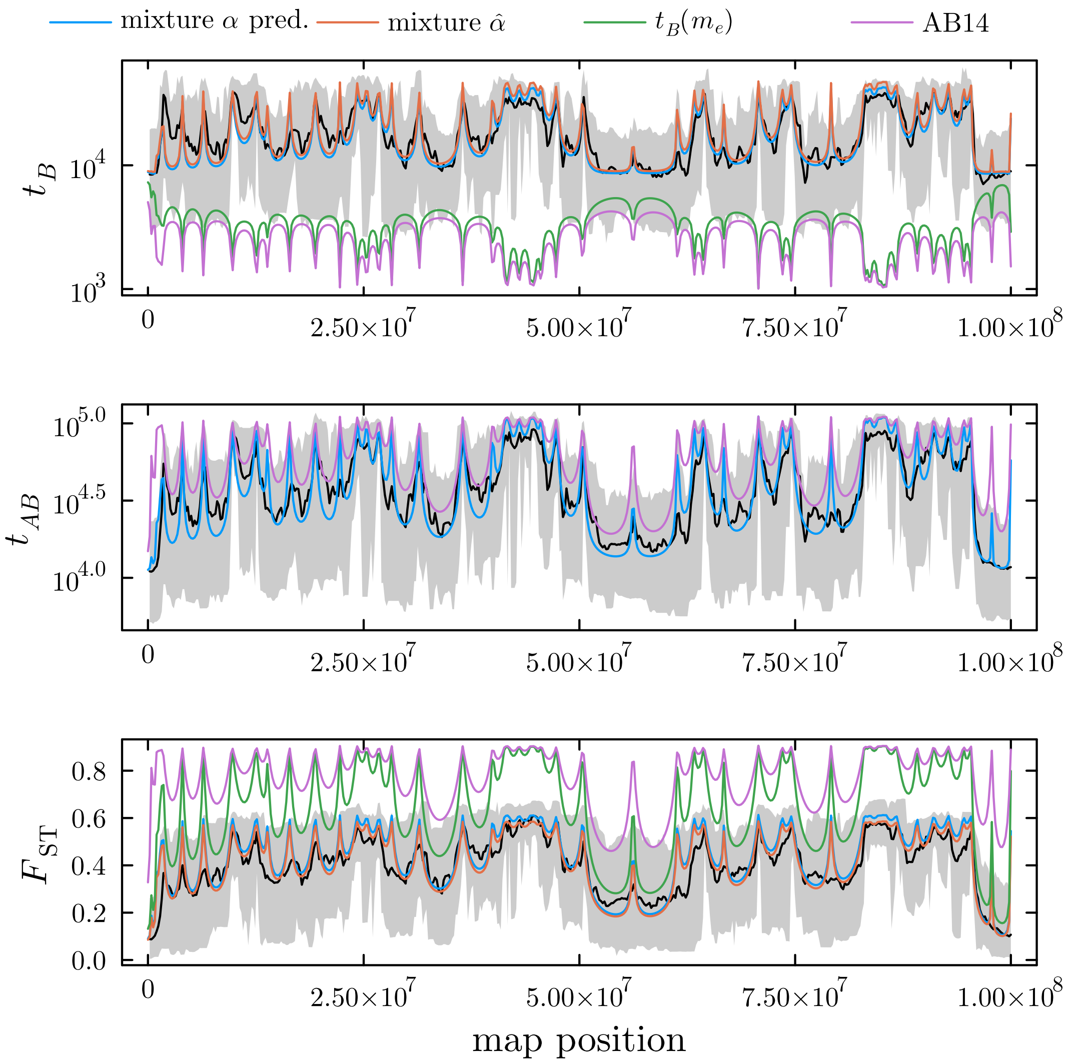

**Figure S21:** As in fig. S19 but with parameters: 10cM, $L\bar{s} = 0.25, L = 50, N_B\bar{s} = 5, m/\bar{s} = 8, \hat{\alpha} = 0.44$.

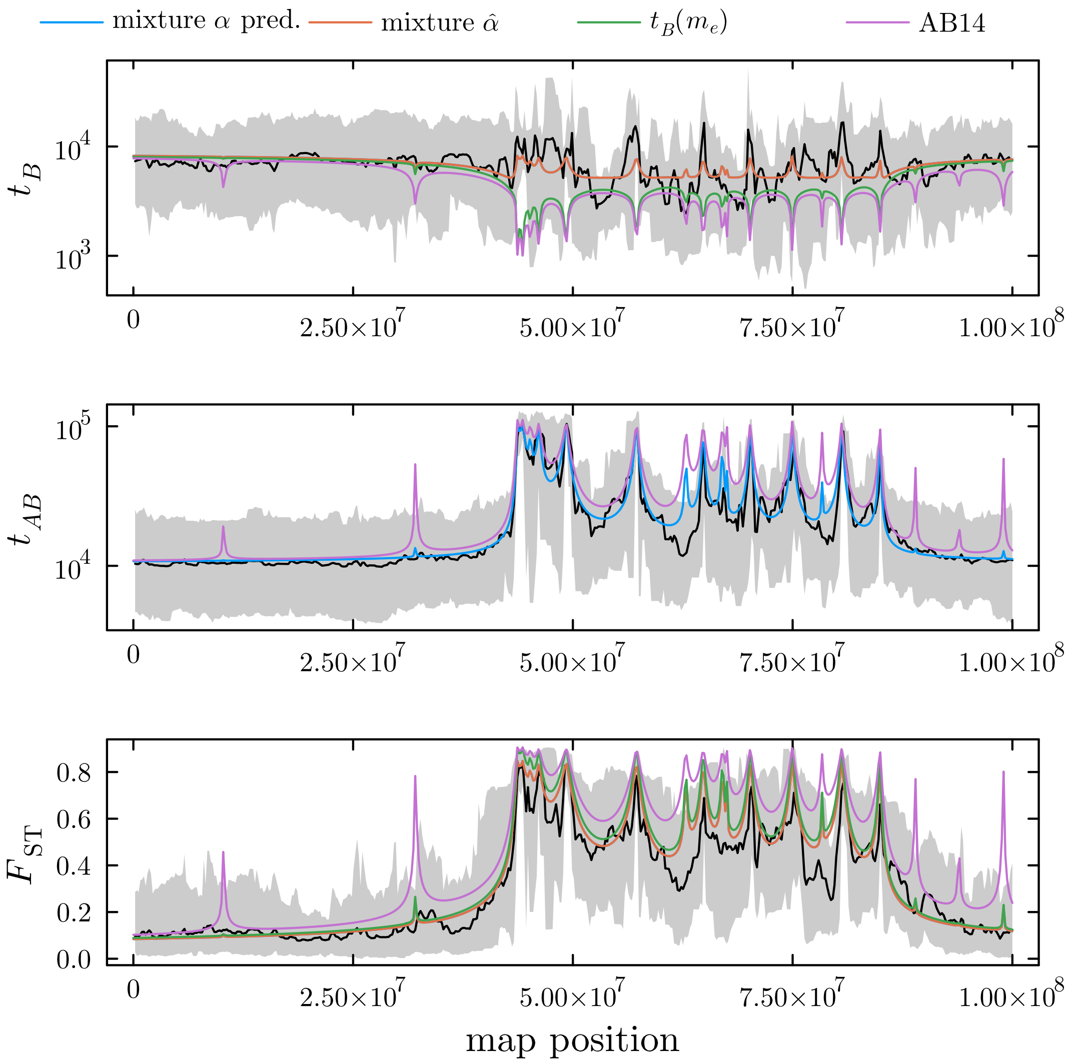

**Figure S22:** As in fig. S19 but with parameters: 10cM, $L\bar{s} = 0.1, L = 20, N_B\bar{s} = 5, m/\bar{s} = 0.5, \hat{\alpha} = 0.07$.

### References

S. Aeschbacher and R. Bürger. The effect of linkage on establishment and survival of locally beneficial mutations. *Genetics*, 197(1):317–336, 2014.

R. Bürger and A. Akerman. The effects of linkage and gene flow on local adaptation: a two-locus continent–island model. *Theoretical population biology*, 80(4):272–288, 2011.

P. Haccou, P. Jagers, and V. A. Vatutin. *Branching processes: variation, growth, and extinction of populations*. Number 5. Cambridge university press, 2005.

M. Nordborg. Structured coalescent processes on different time scales. *Genetics*, 146(4):1501–1514, 1997.

D. Petry. The effect on neutral gene flow of selection at a linked locus. *Theoretical population biology*, 23(3):300–313, 1983.

C. Veller, N. B. Edelman, P. Muralidhar, and M. A. Nowak. Recombination and selection against introgressed dna. *Evolution*, 77(4):1131–1144, 2023.
